# Revising the global biogeography of plant life cycles

**DOI:** 10.1101/2022.09.13.507878

**Authors:** Tyler Poppenwimer, Itay Mayrose, Niv DeMalach

**Affiliations:** School of Plant Sciences and Food Security, Tel Aviv University; Tel Aviv, 6997801, Israel; Institute of Plant Sciences and Genetics in Agriculture, The Hebrew University of Jerusalem; Rehovot, 7610001, Israel

## Abstract

Plants exhibit two primary life cycles – annual and perennial – which vary in their effects on ecosystem functioning. Here, we assembled a database of 235,000 species to assess the worldwide distribution of plant life cycles. We found that annuals are half as common as previously thought (6% of all plant species). Furthermore, our analysis demonstrates that annuals are favored under hot and dry conditions, especially under a prolonged dry season. Strikingly, this pattern remains consistent among different families, indicating convergent evolution. Moreover, we show that increasing climate variability and anthropogenic disturbance further increase the favorability of annuals. Overall, our analysis raises concerns for the future of ecosystem services provided by perennials because the ongoing climate and land-use changes are leading to an annuals-dominated world.

**One-Sentence Summary:** This extensive update to plant life cycle biogeography deciphers their dependence on temperature, rainfall, and disturbance.

## Introduction

At its broadest scale, terrestrial plants can be categorized into two main types of life cycles, annual and perennial (*1, 2*). These growth forms represent the most fundamental characteristic of plant species and illustrate the inherent trade-offs between reproduction, survival, and seedling success (*1, 3*). Annual plants reproduce once and complete their life cycle within one growing season, while perennial plants live for many years and, in most cases, reproduce multiple times. The evolutionary trade-offs reflected in these strategies manifest in numerous functional attributes, such as leaf (*4*) and root traits (*5*), invasiveness (*6, 7*), genome characteristics (*8*), and community stability (*9*) and therefore have many consequences for ecosystem functioning and services (*10, 11*). For example, by allocating more resources belowground, perennials reduce erosion, store organic carbon, and have higher nutrient- and water-use efficiencies (*11–14*).

The differences between annual and perennial plants are noticeably reflected in agricultural settings. Despite being a minor part of global biomass (*15*), annual species are the primary food source of humankind, probably because they allocate more resources to seed output, thereby enhancing agricultural productivity. Annual plants cover 70% of the croplands and provide 80% of worldwide food consumption (*16*). During the Anthropocene, the global cover of annuals dramatically increased because natural systems, often dominated by perennials, were converted into annual cropland (*17, 18*). Moreover, the proportion of annuals increases in many systems because woody perennials have a higher extinction rate (*19*), while invasive species tend to be annuals (*6*).

The annual life cycle has repeatedly evolved in different families, suggesting that it provides a fitness advantage under certain conditions (*20*). The classical theory proposes that the prevalence of life cycle strategies is driven by the survivorship ratio between seeds and adults (*21, 22*). Accordingly, annual species should be more favored under arid conditions that may constrain the persistence of adults. In contrast, wetter conditions, which increase adult persistence, should increase perennial species’ favorability (*23*). Similar arguments can also be made for climate unpredictability and anthropogenic disturbance that strengthen the selection for the annual life cycle (*1, 20, 24–27*).

Numerous studies have discussed plant life cycles as primary examples of adaptation to different climatic conditions and provide estimates for their prevalence in various regions (*28*– *31*). However, the data provided in many of these studies, which penetrated many current ecological textbooks (*30, 31*), are problematic in several aspects. First, the current estimate for the global proportion of annual species (13%) is based on a century-old sample of merely 400 species (*2*). Second, current biome level estimates are based on a single location and extrapolated to represent the entire biome. For example, the desert biome is assumed to contain 42% annual species (*29, 30*) but is based only on data from the Death Valley in California (*2*). Third, current estimates are inconsistent and difficult to compare due to ambiguous biome definitions. For example, an alternative estimate for the desert biome suggests that 73% of plant species are annuals (*28, 31*). Lastly, each biome incorporates a wide range of conditions, e.g., the mean temperature in the desert biome ranges from 30° to −10°C. Thus, this definition aggregates regions that differ markedly in their environmental conditions, likely affecting the prevalence of the different life cycles.

As central as plant life cycles are to plant ecology and evolutionary research, it is remarkable that we still have no clear estimate for the worldwide prevalence of life cycles and their environmental drivers. Yet, such an assessment is essential in times of climate and land-use changes (*17, 32*), which are expected to dramatically alter patterns of plant biogeography with many consequences for ecosystem processes and services (*33–36*). Here, we present the largest assemblage of plant life cycle data encompassing over 235,000 plant species. We cross this database with millions of georeferenced data points. This enormous dataset enables us to revise the biogeography of plant life cycles and showcase the first worldwide map of their distribution. Furthermore, it allows us to evaluate key ecological hypotheses for plant life cycle strategies that were never tested on a global scale. Inspired by ecological theory that predicts annual species to be favored with increasing adult mortality (*21, 22*), we expected the proportion of annuals to increase in conditions favoring adult mortality, including: (1) low precipitation and high temperature, (2) high interannual rainfall variability, and (3) high disturbance.

## Results

We have assembled an extensive plant growth form database containing life cycle data for ≈ 67% of all vascular plant species and georeferenced data for ≈ 51%. This compilation revealed dramatic differences in the relative prevalence of annual and perennial plant species compared to existing estimates. Specifically, the annual life cycle is half as common as previously thought (*28–31*), with annual species comprising ≈6% of all species and ≈15% of herbaceous species (i.e., omitting all woody species, which are all perennial). Furthermore, our biome-level estimates are also very different than those found in current textbooks (Table 1). For example, in the desert biome annuals are 14% of all species and 25% of herbs, much less than the 42% and 63% reported previously, while in the Tundra, the proportion of annuals is much higher than the previous estimates (See Table S31 for a comparison with another old estimate found in (*31*)). Below, we focus on the proportion of annual species among herbaceous species (rather than among all species), which provides a better ‘resolution’ in regions with a high proportion of woody species. Nonetheless, the same results were obtained when we analyzed the proportion of annuals among all species (Figs. S6-S10 and Tables S15-S28).

**Table 1.**
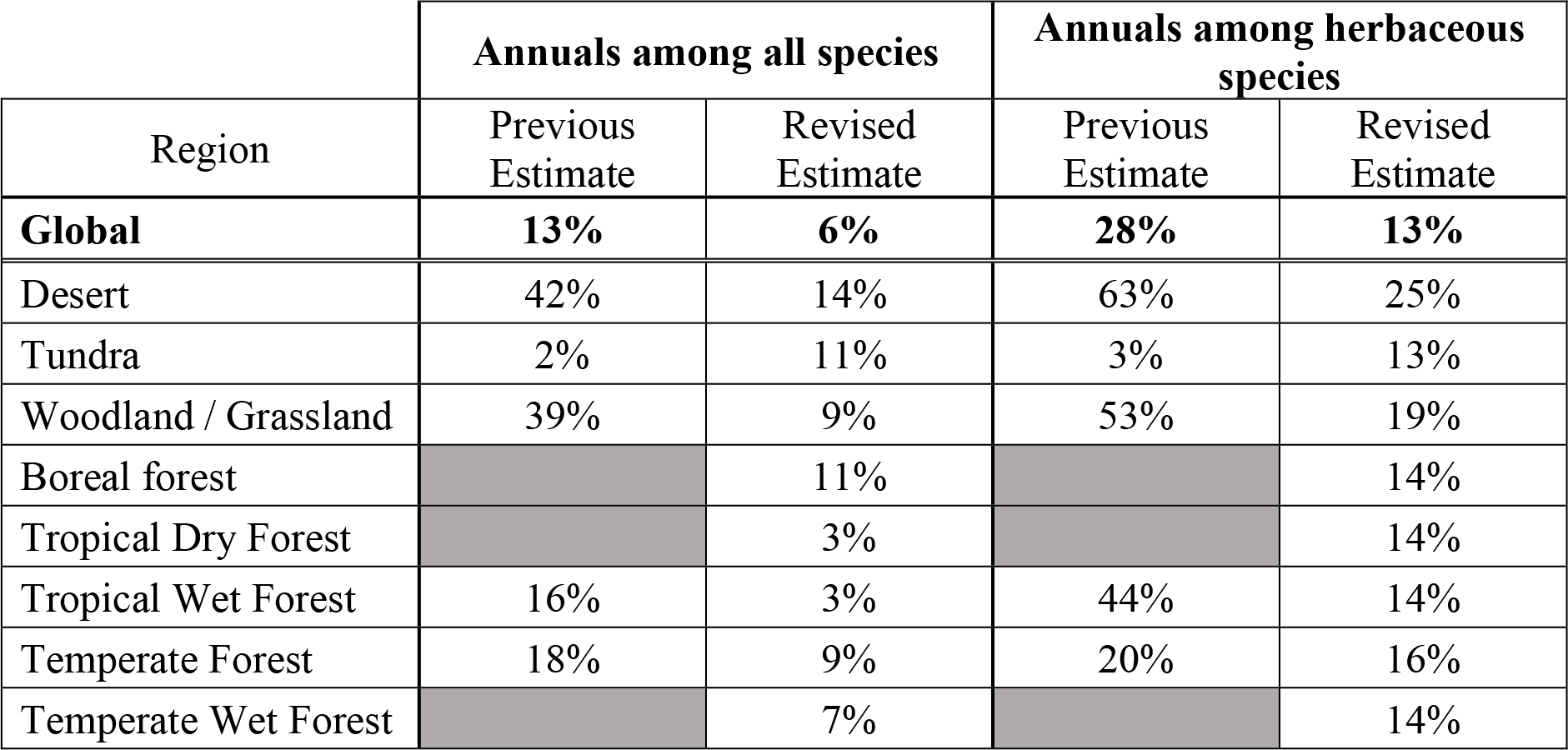
A comparison of previous estimates, obtained from (*30*), for the proportion of annuals among all species and among herbaceous species to our revised estimates. Greyed cells have no initial biome estimate. Alternative previous estimates are available in Table S31. Note, the biome nomenclature used for the previous estimates differs from ours and so the location of the original study was used to determine the corresponding biome. Additional information can be found in Table S30.

| Region | Annuals among all species |  | Annuals among herbaceous species |  |
| --- | --- | --- | --- | --- |
|  | Previous Estimate | Revised Estimate | Previous Estimate | Revised Estimate |
| **Global** | **13%** | **6%** | **28%** | **13%** |
| Desert | 42% | 14% | 63% | 25% |
| Tundra | 2% | 11% | 3% | 13% |
| Woodland / Grassland | 39% | 9% | 53% | 19% |
| Boreal forest |  | 11% |  | 14% |
| Tropical Dry Forest |  | 3% |  | 14% |
| Tropical Wet Forest | 16% | 3% | 44% | 14% |
| Temperate Forest | 18% | 9% | 20% | 16% |
| Temperate Wet Forest |  | 7% |  | 14% |

Overall, we found that ecoregions dominated by annuals are rare, with only 5.5% of ecoregions exhibiting an annual herbs proportion of 50% or more (Fig. 1). Most of these ecoregions were located in regions with desert/semi-desert and Mediterranean climates. In accordance, at the biome level (following Whittaker’s approach (*37*)), we supported the predictions that annuals are favored with increasing temperature and lower precipitation (Fig. 2A). Still, the differences among biomes were not substantial, with the proportion of annual herbs ranging from 13% to 25%, suggesting that the role of climate is underestimated in this coarse spatial scale.

**Fig. 1.**
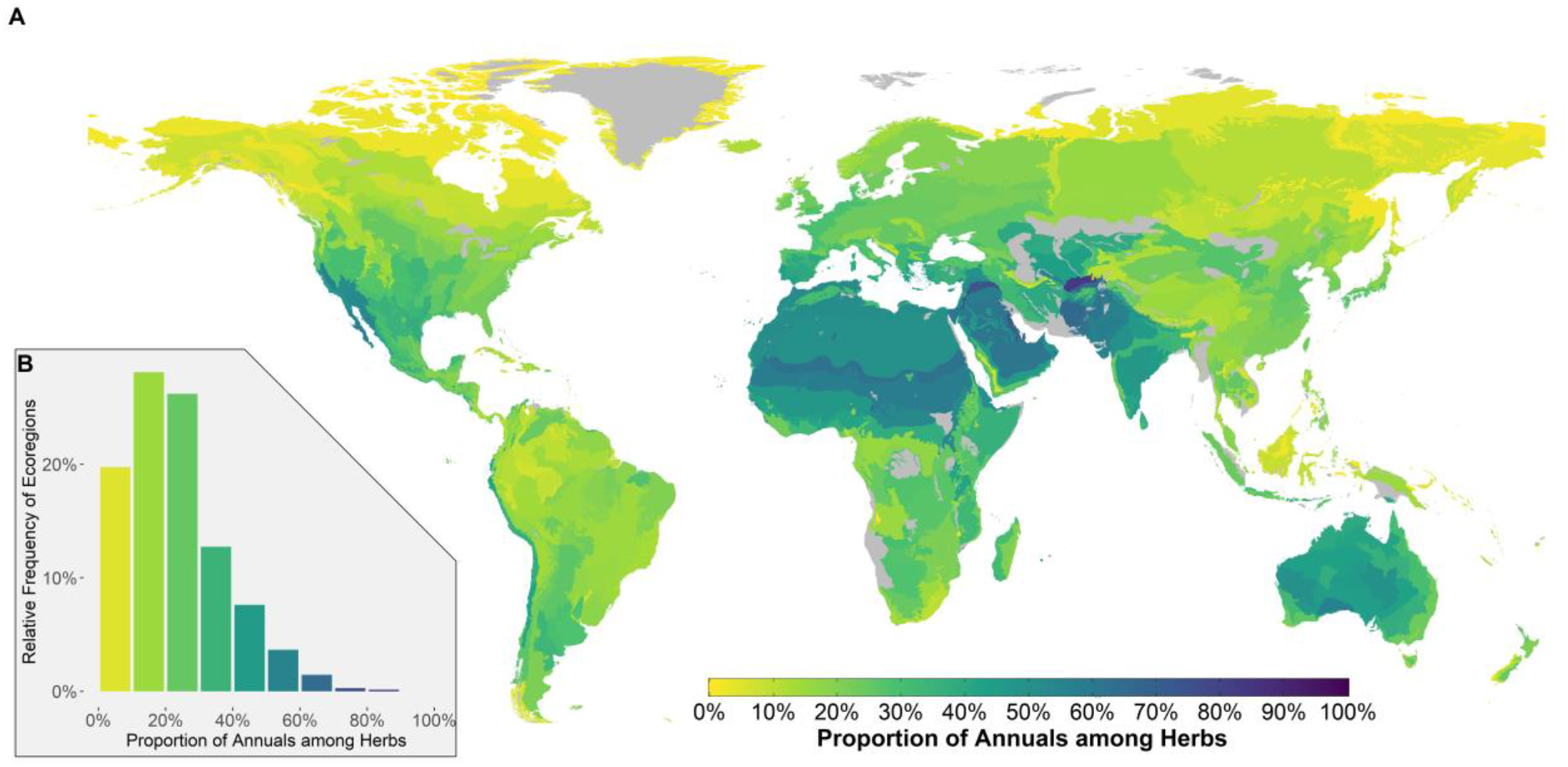
The worldwide biogeography of annuals prevalence. (**A**) A map of the proportion of annual species (among herbaceous species) in each ecoregion. (**B**) The distribution of annuals proportions among ecoregions. Ecoregions with insufficient data (see Methods) are colored grey resulting in 683 colored ecoregions.

**Fig. 2.**
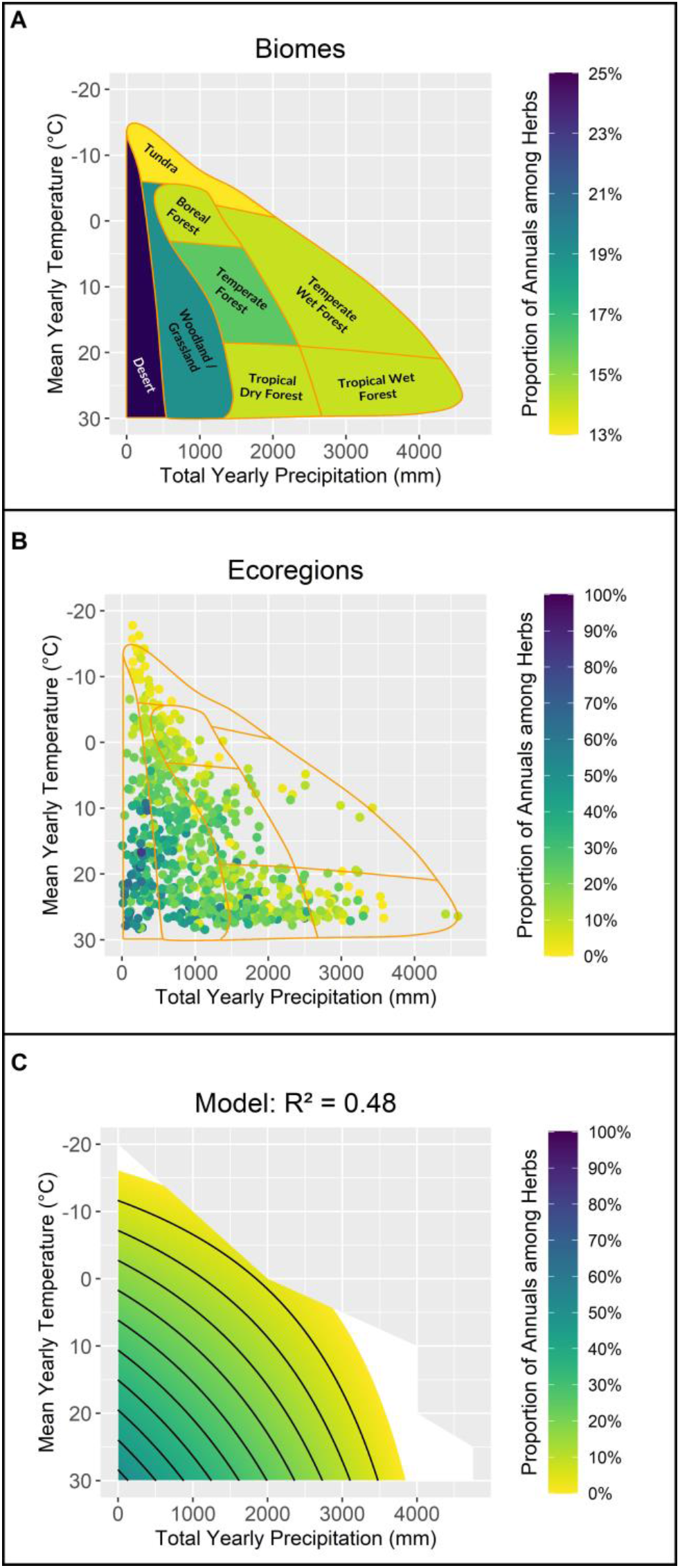
The effects of total yearly precipitation and mean yearly temperature on the proportion of annuals (among herbaceous species). (**A**) The proportion of annuals in each of Whittaker’s biomes. (**B**) The proportion of annuals in each ecoregion (the outline of Whittaker’s biomes is marked by orange lines). (**C**) Predictions of a model of the proportion of annuals as a function of the mean yearly precipitation and temperature (contour lines every 5%). Note that the scale is different for panel (A). N=683.

The large variability within each biome can be seen when examining the proportion of annuals at the ecoregion resolution (Fig. 2B). For example, not all desert-biome ecoregions have a high proportion of annual herb species, with cooler deserts exhibiting much lower proportions than hot ones. The same trend is repeated among other biomes, as ecoregions with lower precipitation and hotter temperatures (i.e., located in the lower-left coordinate of their biome in Fig. 2B) possess greater annuals. This pattern was corroborated using a linear model that fitted the proportion of annuals as a function of *mean yearly temperature* and *total yearly precipitation*. Notably, these two climatic variables accounted for nearly half of the variance of the worldwide distribution of plant life cycle strategies (*P* <10^−15^, *D. F*. = 679, R^2^ = 0.48), providing global support for the hypothesis that annual plants would be more common under hotter and drier conditions.

The strong relationship between life cycle and aridity could result from phylogenetic artifacts wherein taxonomic groups adapted to arid conditions also have a high proportion of annual species. Assuming the annual life cycle strategy is a general adaptation to hot and dry conditions that had repeatedly occurred in multiple clades (rather than a correlated trait), we expected the proportion to increase under hot and dry conditions within each plant family. We tested this prediction by fitting the same linear model on the four most annual-rich families (Asteraceae, Brassicaceae, Fabaceae, and Poaceae). Strikingly, the same relationship was obtained among the four families, with an explained variation of annual herb proportions ranging from 68% in Asteraceae to 32% in Fabaceae (in all models *P* <10^−15^; Fig. 3).

**Fig. 3.**
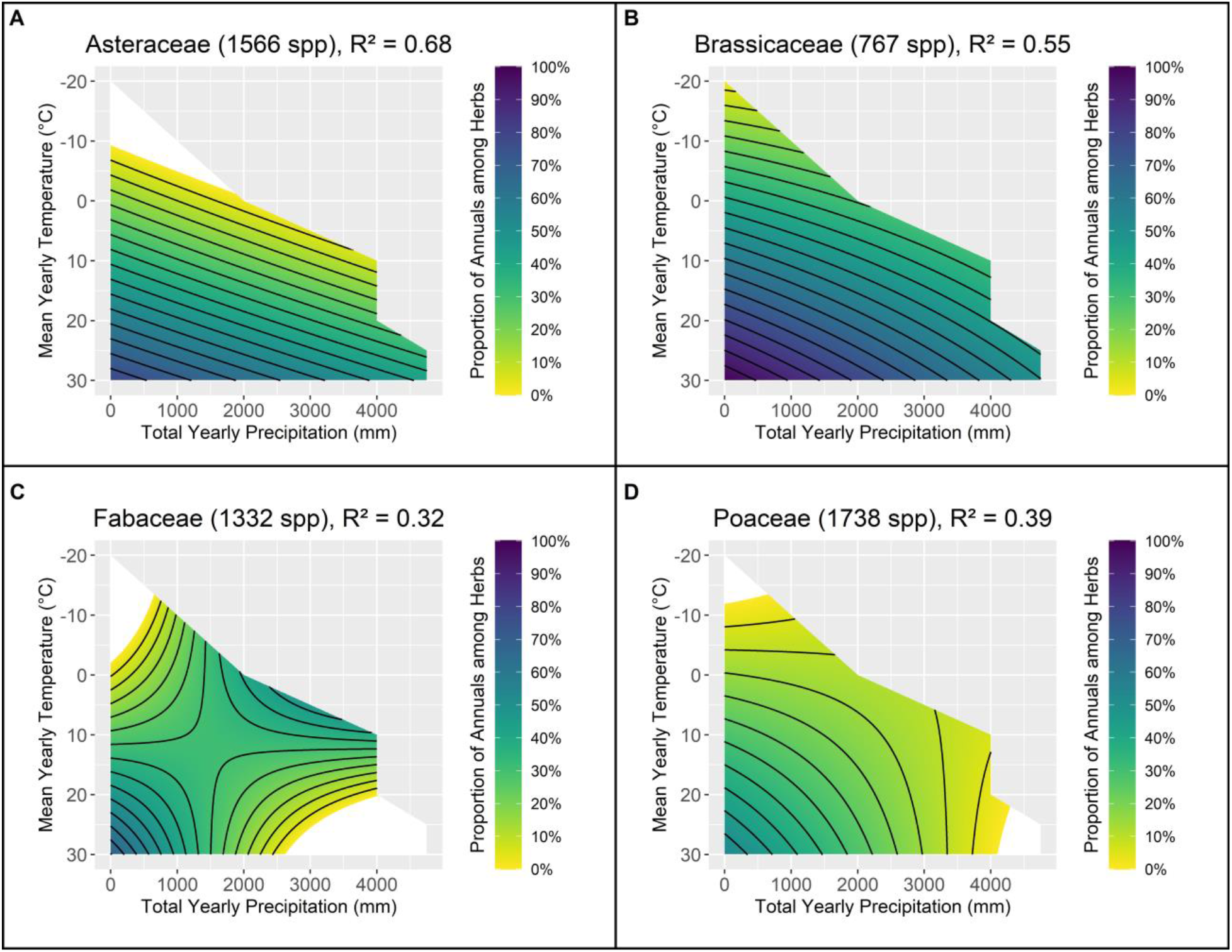
The effects of total yearly precipitation and mean yearly temperature on the proportion of annuals (among herbaceous species) in the four most annual-rich families (predictions of the regression model). (**A**) Asteraceae, N = 465. (**B**) Brassicaceae N = 262. (**C**) Fabaceae N = 382. (**D**) Poaceae N = 513. Contour lines are drawn every 5% for all figures.

Although the model of yearly temperature and precipitation provides a good description of annual herbs proportions across the globe, it does not account for temporal variation in climate throughout the year. We, therefore, fitted a set of two-variable regression models. Each model consisted of one *quarterly* temperature variable and one quarterly precipitation variable. The best-fit model (hereafter the temporal-inclusion model) incorporated the *mean temperature of the warmest quarter* and the log-transformed *precipitation of the warmest quarter* and accounted for 55% of the observed variance (*P* <10^−15^, *D. F*. = 679). According to this model, annual herbs proportion increases with increasing temperature and decreasing precipitation during the warmest quarter (Fig. 4). Furthermore, this model had a substantially better fit than the model based on the *mean yearly temperature* and *total yearly precipitation* outperforming it in terms of explained variance (0.55 vs. 0.48) and information theory criteria (ΔAICc = 92.4).

**Fig. 4.**
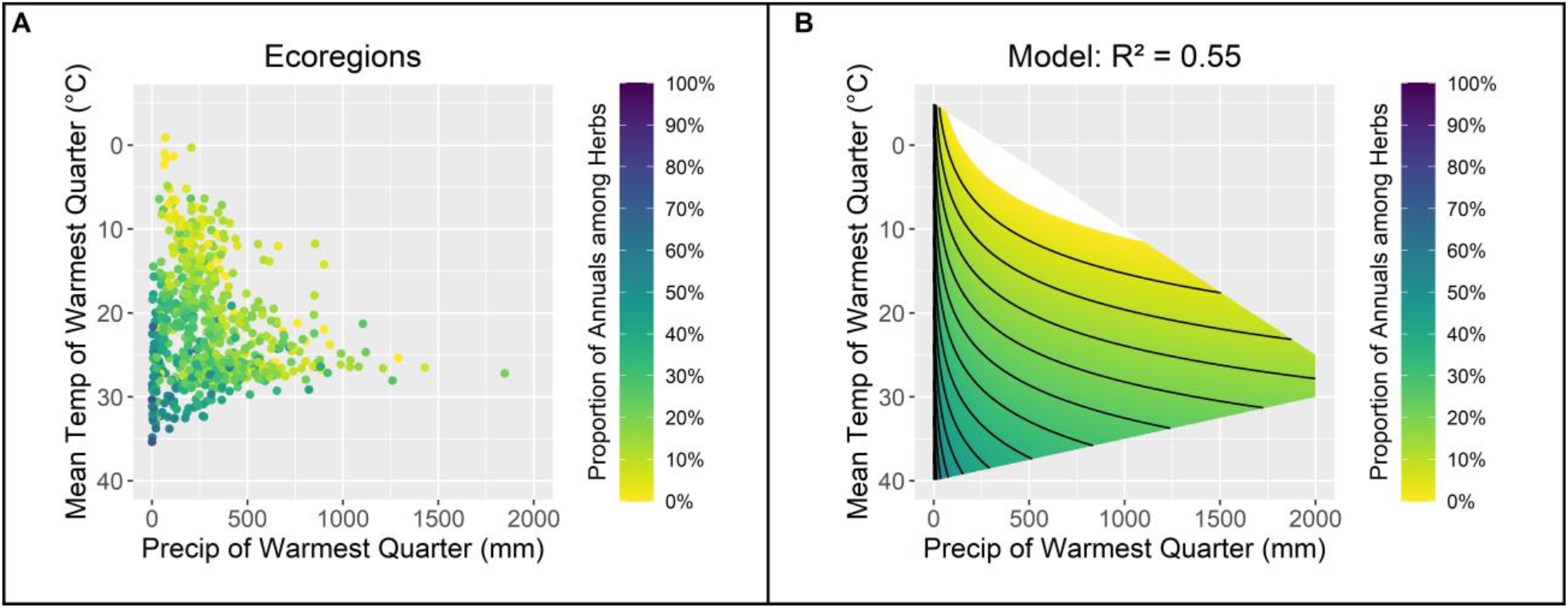
The effects of the total precipitation and mean temperature of the warmest quarter on on the proportion of annuals (among herbaceous species). (**A**) The proportion of annuals in each ecoregion. (**B**) The predictions of a regression model of annual proportion as a function of precipitation and mean temperature of the warmest quarter with contour lines every 5%. N=683.

The quarterly temperature and precipitation model can distinguish between ecoregions with similar annual climate patterns yet a different proportion of annuals. For example, the Eastern Mediterranean (e.g., Tel Aviv) and Chihuahuan desert (southwestern United States and northern Mexico) ecoregions have identical mean yearly temperatures (17.6° C) and relatively similar amounts of yearly precipitation (527mm and 330mm) yet maintain different annual herbs proportion (51% and 36%). Given their similar mean yearly climate, the yearly temperature and precipitation model predicts similar annual herbs proportion for these two ecoregions (32% and 34%, respectively). However, the Tel Aviv ecoregion receives substantially less precipitation (6mm) than the Chihuahuan desert (157mm) during the hottest quarter. As such, the quarterly temperature and precipitation model better differentiates the two ecoregions, producing substantially better predictions (annual herbs proportion of 52% in Tel Aviv and 29% in the Chihuahuan desert).

After accounting for variability within each year, we tested whether accounting for *interannual* variation in precipitation (a proxy for climate unpredictability) can further improve the model. In accordance with the classical theory, we found that increasing climate unpredictability is associated with higher proportion of annual species (*P* <10^−15^, *D*.*F*. = 680, *R*^2^ = 0.243) and further improves the fit of the model that also included quarterly temperature and precipitation (ΔAICc = 45, change in *R*^2^ from 0.55 to 0.58).

Lastly, we tested the prediction that disturbance (human footprint) will increase the proportion of annuals. Indeed, there was a positive effect of the human footprint on the proportion of annuals (*P* < 10^−7^, *D*.*F*. = 681, *R*^2^ = 0.046). Notably, despite the mild individual explanatory power of human footprint, adding this variable to the model with climate unpredictability and quarterly temperature and precipitation further improved the explanatory power of the model (ΔAICc = 24, change in *R*^2^ from 0.58 to 0.61), suggesting that the human footprint index captures patterns the climatic features do not.

## Discussion

This study provides an extensive update to the worldwide biogeography of plant life cycles. We have focused on the proportion of annual species rather than on their relative abundance. Hence, our analysis should primarily reflect the outcomes of evolutionary and biogeographical processes, such as speciation, extinction, and long-scale dispersal, rather than purely ecological processes. Our analysis indicates that annual species are half as common as previously thought (*28–31*). Similarly, our biome-level estimates vary markedly from earlier estimates (e.g., reducing by 3-5 fold the estimated proportion of annual species among desert plants) (Table 1 and Table S31).

Our analysis supports the classical hypothesis that the annual life cycle is favored under hotter and drier conditions. However, yearly means provide an insufficient explanation for some observed patterns, e.g., for the finding that some Mediterranean regions exhibit higher proportions of annual species than hot deserts, despite being, on average, cooler and wetter. We discovered that a long dry summer is a principal factor governing the occurrence of annual-rich regions, demonstrating that the temporal distribution of hot and dry periods is more important than having an arid climate *per se*. We speculate that dry summer conditions result in either lower adult survival or higher seed survival. Both align with established theory that predicts annuals to be favored as the ratio between seedlings/seed survival and adult survival increases (*21, 22*).

Our work also demonstrates that annual prevalence is not only affected by climate patterns. We found that human-mediated disturbance increases the favorability of annual plants. When examined independently, anthropogenic disturbance explains 4.6% of the variability in annual herb proportions. However, its incorporation into the combined model with temperature, precipitation, and climate unpredictability attributes led to a 3% increase in model explanatory power, indicating that a major fraction of the effect of human disturbance is independent of climatic patterns. Bearing in mind that humans have only been present for 300-400 thousand years, a minute in evolutionary time scales and that the effect of human disturbance is exceptionally biased towards recent years, this effect should not be treated lightly.

Overall, our findings raise concerns for the future of perennial species, which often provide more ecosystem services than annual species (*4-14*). First, in line with the projected climate change regime, the anticipated increase in temperatures will drive worldwide conditions towards those that favor annual species. Second, reduced precipitation and the shortening of rainy seasons will prolong low-precipitation summer conditions and increase drought events. Third, the expected intensification of inter-annual variability will result in increasingly unpredictable climate patterns, further favoring annual plants. Finally, with the human population predicted to reach 11 billion by 2100, the impact of anthropogenic activities is expected to play an increasing role in shaping patterns of plant biogeography. Consequently, the current projections of many global conditions are becoming more and more favorable for annual species, ultimately leading to a future world that is increasingly annuals dominated.

## Acknowledgments

We thank Anna Rice for data processing support and Ron Milo for manuscript critique and feedback.

## Funding

Edmond J. Safra Center for Bioinformatics at Tel-Aviv University

## Author contributions

Conceptualization: TP, IM, ND

Data Curation: TP

Formal Analysis: TP

Funding Acquisition:

Investigation: TP

Methodology: TP

Project Administration: IM, ND

Resources: IM

Software: TP

Supervision: IM, ND

Validation: TP

Visualization: TP

Writing – Original Draft Preparation: TP

Writing – Review & Editing: TP, IM, ND

## Competing interests

Authors declare that they have no competing interests.

## Data and materials availability

All data and code are available at https://doi.org/10.6084/m9.figshare.c.6176239.v1.

## Supplementary Materials

Materials and Methods

Figs. S1 to S10

Tables S1 to S35

References (*38–51*)

## Notes

https://doi.org/10.6084/m9.figshare.c.6176239.v1

### Competing Interest Statement

The authors have declared no competing interest.

