## Supplementary Materials for "Revising the global biogeography of plant life cycles"

Science
AAAS

### Supplementary Materials for

#### Revising the global biogeography of plant life cycles

Tyler Poppenwimer[1,2], Itay Mayrose[1*], Niv DeMalach[2]

**This PDF file includes:**

Materials and Methods
Figs. S1 to S10
Tables S1 to S35

#### Materials and Methods

Life Cycle database development

We built an extensive life-cycle database by aggregating all types of vascular plant data from 11 disparate plant trait databases (*10*, *38-47*) (see Table S35 for access dates). This raw database contained approximately 6.4 million entries and 400,000 unique names. All unique names were resolved using the R package *WorldFloraOnline* v1.7 (WFO) to ensure a uniform naming scheme and exclude unrecognized species. Resolved names were filtered by match distance and WFO acceptance (see Supplementary Text for a full description).

All unique terms were manually assessed to extract data relevant to a plant's life cycle – annual/perennial – and growth form– woody/herbaceous when available. Those that did not provide relevant information or provided conflicting information were excluded. After term interpretation, there were 5.6 million entries and 262,000 unique *species* remaining.

Life cycle consensus among each species' data was achieved by comparing all life cycle and biomass composition entries. Only those species with a unanimous term agreement and without conflicting life cycle and biomass composition consensuses were considered. Crop species were excluded as occurrence data may not represent natural habitats. A list of crop species was obtained from (*10*). This process produced a database of approximately 235,000 species with life cycle information. Our database contains approximately 67% of all WFO accepted plant species names and represents the largest plant life cycle database developed to date.

Matching life cycle data with species observations

Species observation data were based on occurrence data available through the Geographic Biodiversity Information Facility (GBIF) (*48*). All observation data points within the Plantae kingdom (approximately 355 million) were downloaded (September 14$^{th}$, 2021) and processed locally. We filter unreliable data points following the recommendation of the vignette of the R package *CoordinateCleaner* v2.0-18. The following steps were used to filter unreliable data points:

1) Data points without coordinates were excluded
2) The R package *CoordinateCleaner* v2.0-18 was used to discard data points with erroneous locations and problematic temporal metadata (see Supplementary Text for a full description of this process).
3) Data points were removed if the recorded 'coordinate uncertainty' was greater than 100km.
4) Data points whose 'Basis of records' was literature or living specimen were discarded (these generally refer to the location of museum or herbaria collections)

5) Data points whose record date was during or before 1945 were excluded as it has been suggested these may be more unreliable
6) Data points that were not labeled as species

Once cleaned, all remaining unique names were resolved using the WFO package, and the same criteria as in the life form database were applied. Once the names were resolved, the species in the cleaned GBIF database and the assembled life form database were matched. Of the 235,000 species in our assembled lifeform database, approximately 180,000 species were found within the cleaned GBIF data.

To mitigate sampling bias and inexact coordinates, species observation data was mapped into larger geographical regions defined by specific environmental and ecological conditions (*10*). To this end, each georeferenced data point was assigned to one of 827 ecoregions as defined by the World Wildlife Fund (WWF) (*49*). This process was accomplished using the R packages *raster* v3.4-13 and *rgdal* v1.5-27. Following the procedures used by (*10*), species were considered present in a geographic region if there were five or more observations to overcome rare species. Similarly, to ensure all regions contained sufficient data for analysis, each region was only considered if ten or more species were present.

This procedure produced sufficient data for 726 ecoregions when examining annual species among all species and 683 ecoregions for annual species among only herbaceous species.

Predictors of annual proportion

We examined the relationships between various climatic and anthropogenic features and the distribution of plant life cycle strategies. To this end, we determined the frequency of plant life form strategies by considering the number of species with a given trait out of the total number of species with life form data in each region (e.g., annual species out of all species with life cycle data). Each region was subsequently assigned a suite of climatic and anthropogenic features. Unless otherwise indicated, all features were determined by taking the median value across all pixels in a region.

We downloaded bioclimate features from the WorldClim Global Climate Data at ten arc-minutes resolution (*50*). All 19 BIOCLIM variables representing each region's major temperature and precipitation characteristics were extracted using the R package *raster* v3.4-13.

As a measure of climate unpredictability, we calculated interannual precipitation variation by extracting the coefficient of variation (CV) for yearly precipitation from each region using the R package *raster* v3.4-13. We aggregated all available monthly precipitation data layers from the WorldClim Global Climate Data (*50*) at ten arc-minutes resolution (1961 – 2018) to determine the total yearly precipitation for each pixel in each year. The mean and CV of the total yearly precipitation across all years for each pixel were then used to determine the coefficient of variation. Unlike BIOCLIM 3, which determines precipitation variability within a year, our estimate measures precipitation variability between years.

We used the Human Footprint data layer assembled by (*51*) as a proxy for anthropogenic disturbance, which represents the total ecological footprints of the human population. This layer incorporates eight variables: built-up environments, population density, electric power infrastructure, crop lands, pasture lands, roads, railways, and navigable waterways. Together, these features' impact is used to evaluate the amount of land or sea necessary to support human activity's consumption habits. Human Footprint values were extracted for each region using the R package *raster* v3.4-13.

Statistical analyses

In the first step, we regressed annual and annual herb frequency against mean yearly temperature, total yearly precipitation, and their interaction (this model is referred to as the classical model). Then, we applied the same model separately to the four most annual-rich families (*Asteraceae*, *Brassicaceae*, *Fabaceae*, and *Poaceae*).

In the second step, we compared models of two climatic variables using one quarterly temperature bioclimatic variable and one quarterly precipitation variable (this model is referred to as the temporal-inclusion model). We had four temperature bioclimatic features (BIO8, BIO9, BIO10, BIO11) and four precipitation features (BIO16, BIO17, BIO18, BIO19). We omitted month-specific bioclimatic features because they are highly correlated with quarter-specific. Preliminary analysis suggested that log transformations of precipitation bioclimatic features often increased explanatory power, and therefore they were also included in the exhaustive search. Altogether, we had 32 different models (four temperature and eight precipitation features) with an additional two models using mean yearly temperature and total yearly precipitation and the log transformation of total yearly precipitation. Model comparison was achieved using AIC values obtained from the R package *MuMIn* v1.43.17.

In the third step, we investigated the role of CV in annual precipitation (a proxy of climate uncertainty) and human footprint (a proxy for disturbance) on annual and annual herb frequencies. This analysis was conducted on annual species within 725 ecoregions and annuals among herbaceous species within 682 ecoregions. We started by testing each variable alone and then tested whether it increases the fit of the best temporal-inclusion model.

Biome Estimates

To obtain biome estimates of annual and annual herb frequencies, all ecoregions with sufficient data were individually plotted in the total yearly precipitation and mean yearly temperature space with the Whittaker biome overlay outline (adapted from (*37*)) overlaid (see Fig 2B for reference). We determined each ecoregion's biome based on its location within this space. For those ecoregions whose biome designation was difficult to assess, their points were enlarged until one biome had a plurality of the circle's area. For those ecoregions outside Whittaker's biome space, their biome designation was determined by the closest biome. Once the biome designation of all ecoregions was determined, the species presence data for all ecoregions within a given biome were aggregated. The same process used to determine the presence and absence of species in an ecoregion was used to determine the presence and absence of species in the biome. Of note, a biome could have more species than the combined ecoregions within said biome because some species may have five or more observations within the biome, but not within any of the individual ecoregions.

Comparing Previous Biome Estimates

The definition of the biomes used to determine previous biome-level estimates of annuals proportions were not explicitly defined making direct comparison with our set of biomes difficult. However, we traced the origins of each estimate and determined the original study's locations. These locations were then matched with the WWF ecoregions and then the corresponding biome was determined as discussed above. This allowed direct comparison between previous estimates and our revised estimates. Additionally, estimates for the proportion of annuals among herbaceous species was not explicitly provided by previous estimates. For comparison purposes, previous annual herbs proportion estimates were calculated based on the available biome-level life form classification estimates available from each study. See Tables S36 and S38 original study location matchings and annual herbs proportions calculations.

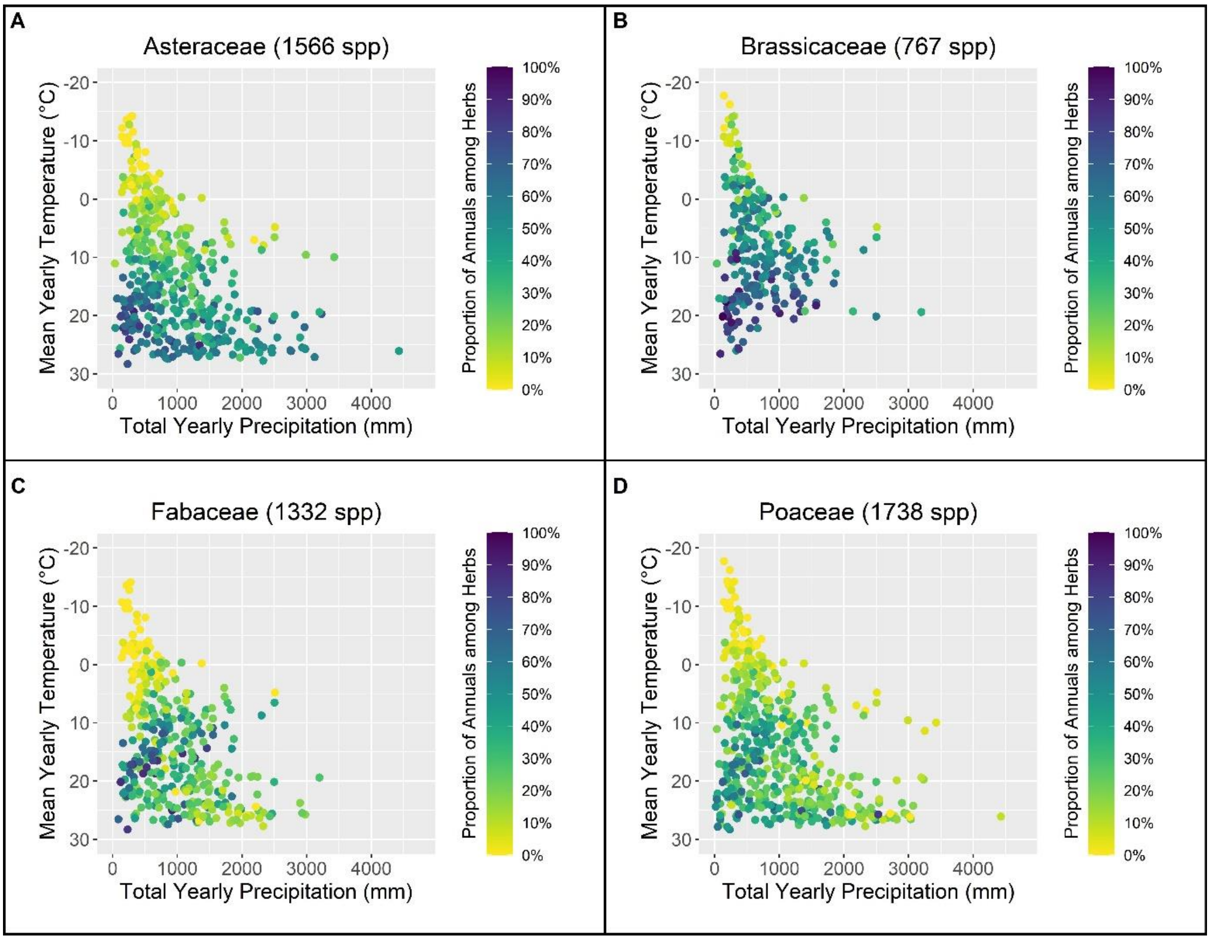


**Fig. S1.**
Scatter plots of the effect of total yearly precipitation and mean yearly temperature on the proportion of annuals among herbs in ecoregions with sufficient data for the four most annual-rich families. (**A**) Asteraceae, N = 510. (**B**) Brassicaceae N = 265. (**C**) Fabaceae N = 527. (**D**) Poaceae N = 517.

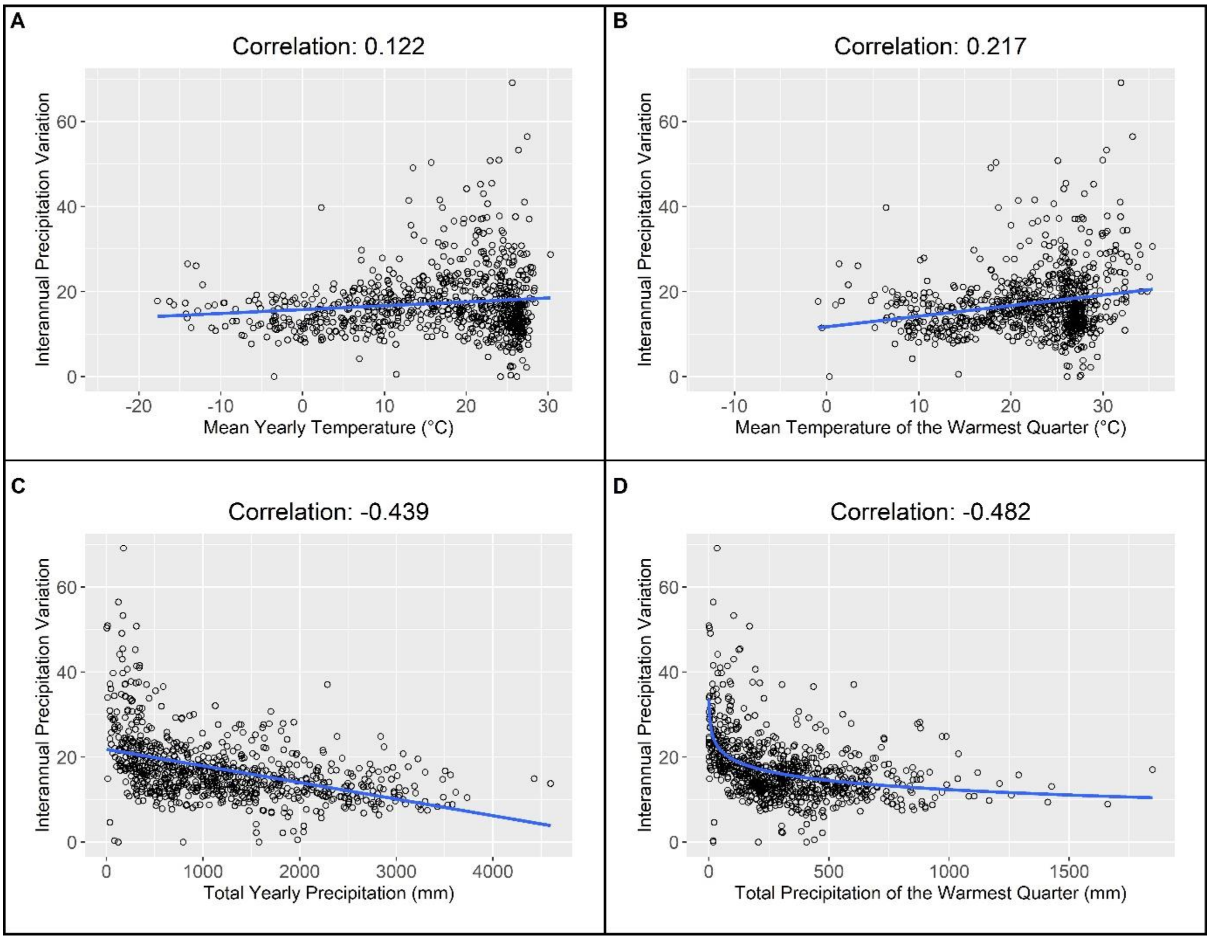


**Fig. S2.**
Scatter plots depicting the correlation between interannual precipitation variation and various BioClim features of each ecoregion with a best fit line (blue) added for easier interpretation. (**A**) BioClim1, mean yearly temperature. (**B**) BioClim10, mean temperature of the warmest quarter. (**C**) BioClim12, total yearly precipitation (**D**) BioClim18, total precipitation of the warmest quarter. Note that correlation and the regression line in (D) are fitted to the log transformation of BioClim18 as used in the quarterly temperature and precipitation model.

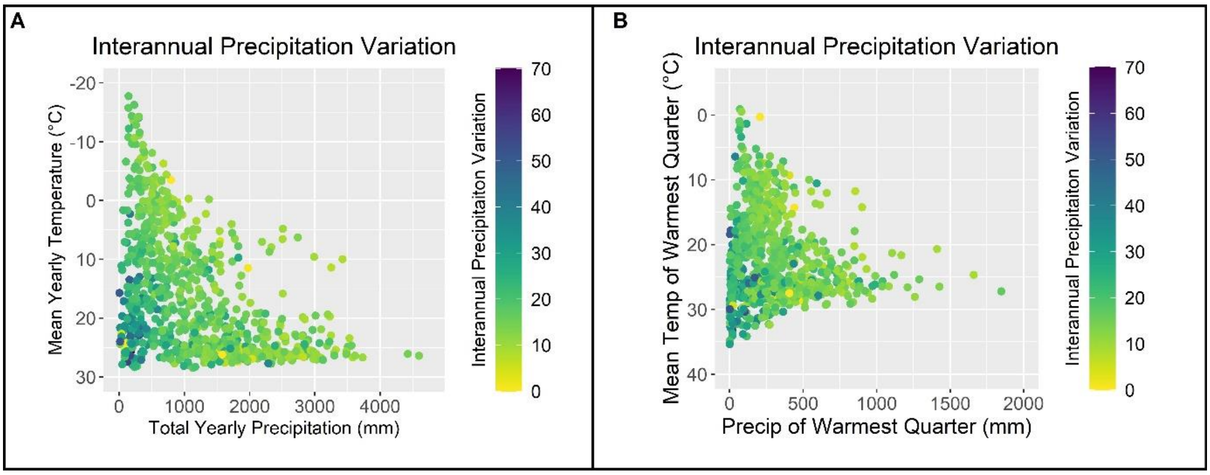


**Fig. S3.**
The relationship between interannual precipitation variation and BioClim feature pairs used in the classical and temporal models in each ecoregion. (**A**) Total yearly precipitation and mean yearly temperature used in the yearly temperature and precipitation model. (**B**) Total precipitation and mean temperature of the warmest quarter use in the quarterly temperature and precipitation model.

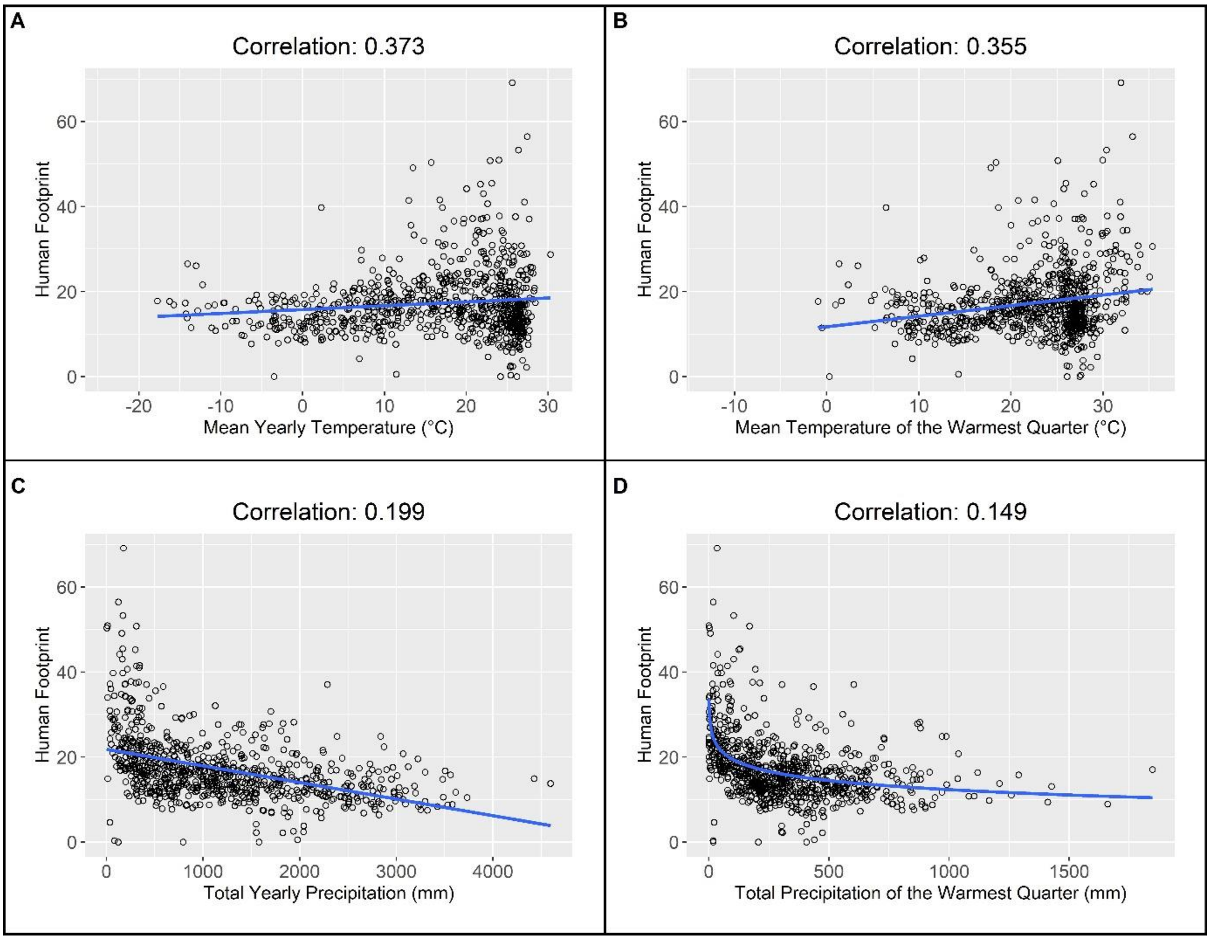


**Fig. S4.**

Scatter plots depicting the correlation between Human Footprint and various BioClim features of each ecoregion with a best fit line (blue) added for easier interpretation. (**A**) BioClim1, mean yearly temperature. (**B**) BioClim10, mean temperature of the warmest quarter. (**C**) BioClim12, total yearly precipitation (**D**) BioClim18, total precipitation of the warmest quarter. Note that correlation and the regression line in (D) are fitted to the log transformation of BioClim18 as used in the quarterly temperature and precipitation model.

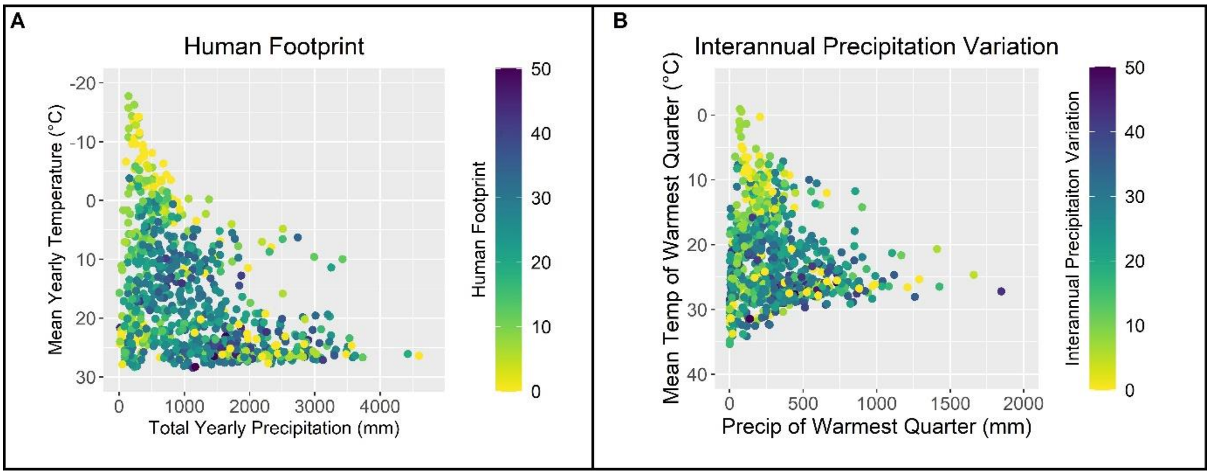


**Fig. S5.**
The relationship between human footprint and BioClim feature pairs used in the classical and temporal models in each ecoregion. (**A**) Total yearly precipitation and mean yearly temperature used in the yearly temperature and precipitation model. (**B**) Total precipitation and mean temperature of the warmest quarter use in the quarterly temperature and precipitation model.

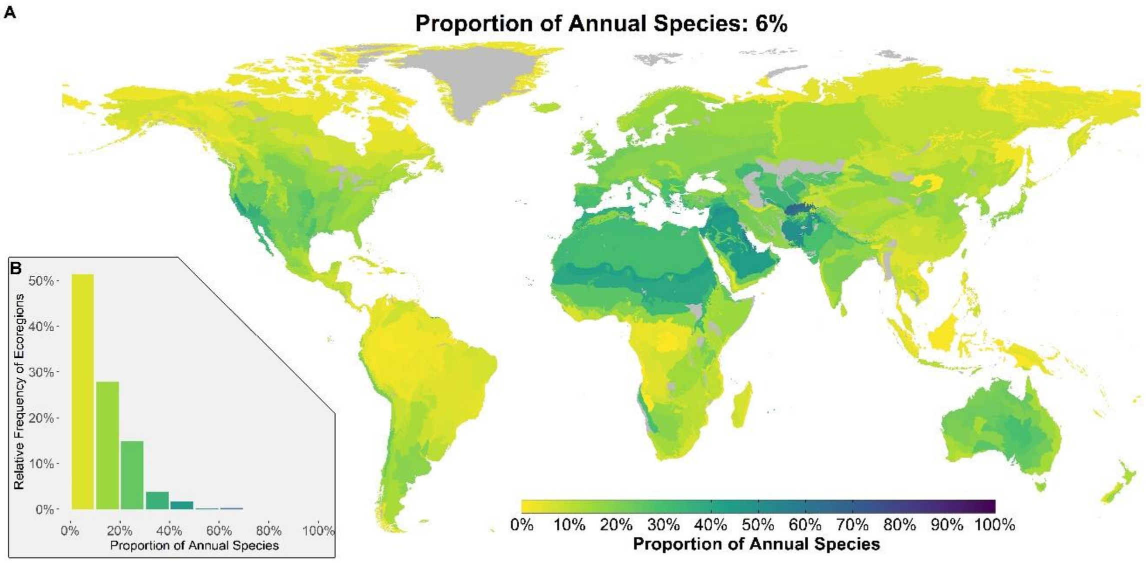


**Fig. S6.**
The worldwide proportion of annuals among all species in ecoregions. (**A**) The proportion of annual species among all species. (**B**) The distribution of annuals proportions among ecoregions. Ecoregions with insufficient data (see Methods) are colored grey resulting in 726 colored ecoregions.

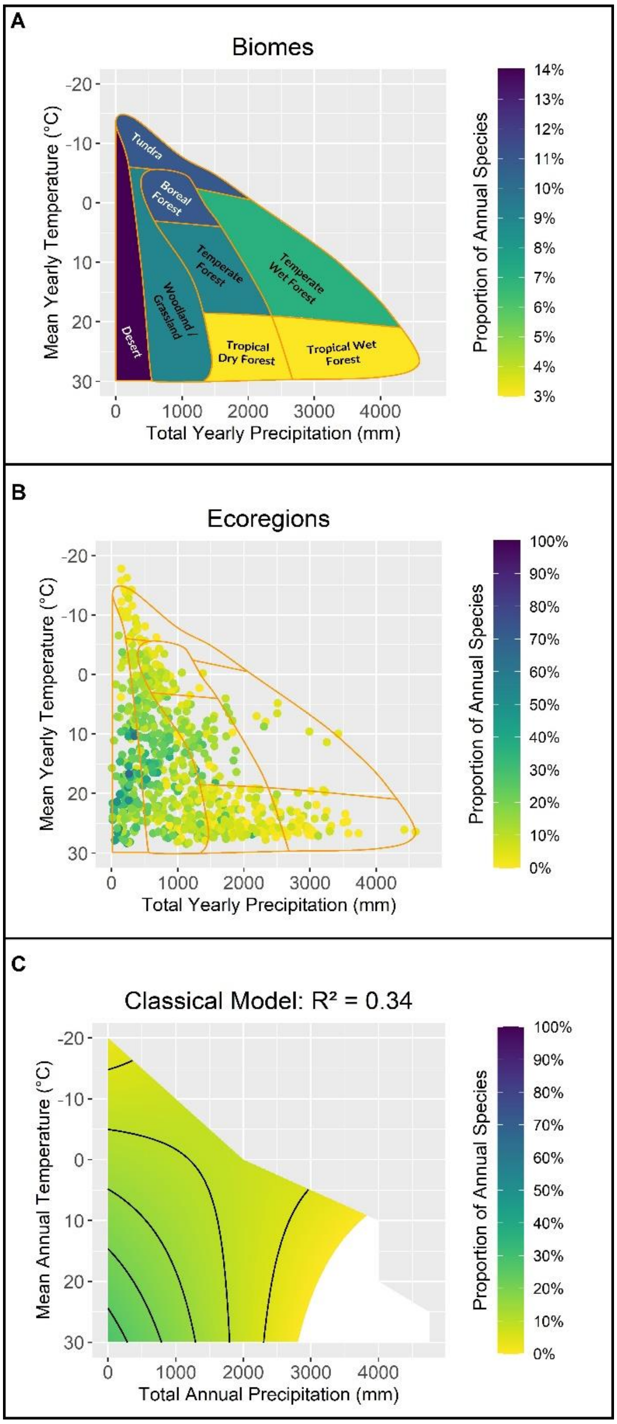


**Fig. S7.**

The effects of mean yearly precipitation and temperature on the proportion of annuals among all species. (**A**) The annuals proportion in each of Whittaker's biomes. (**B**) The annuals proportion in each ecoregion (the outline of Whittaker's biomes is marked by orange lines). (**C**) Predictions of a model of annual frequency as a function of the mean yearly precipitation and temperature (contour lines every 5%). Note that the scale is different for panel (A). N=726.

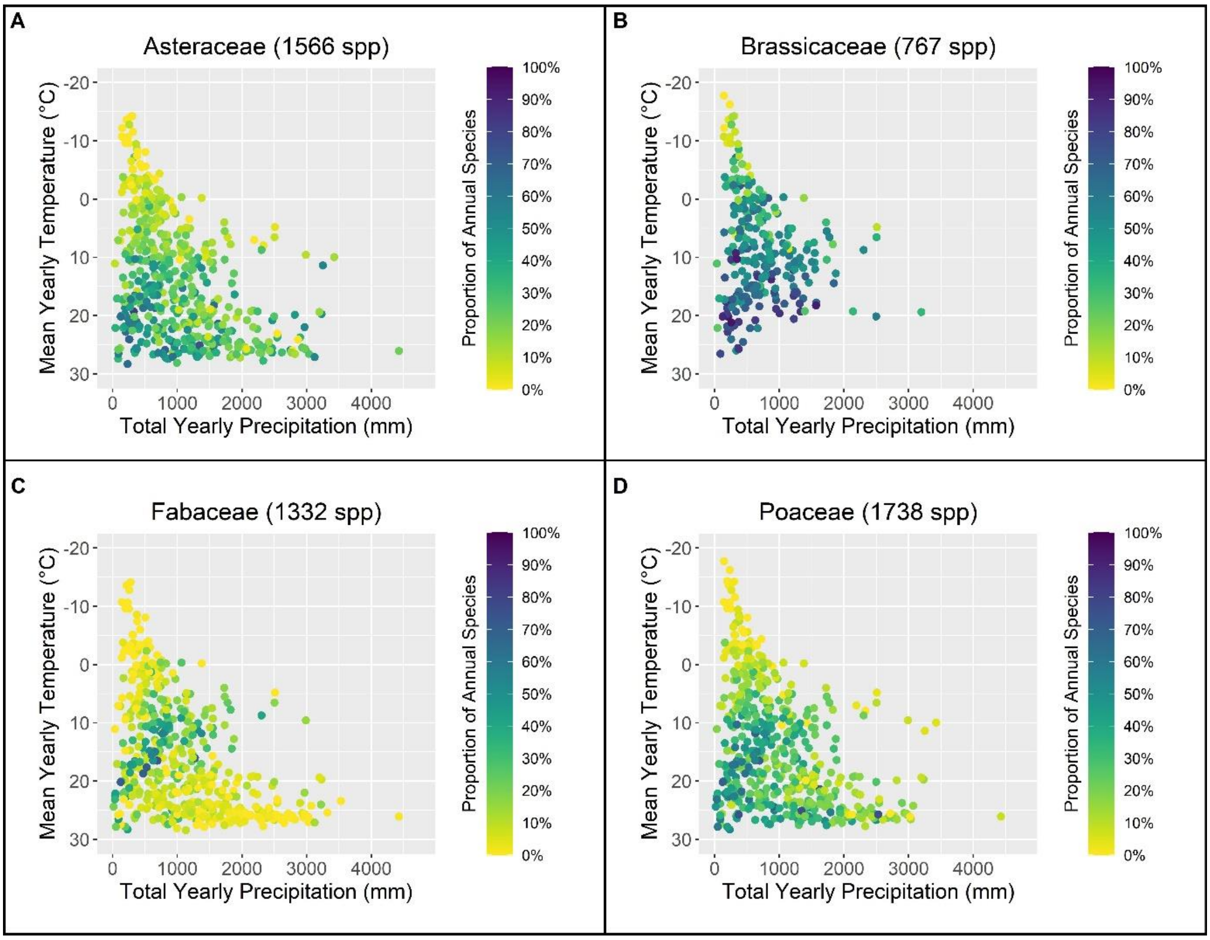


**Fig. S8.**
Scatter plots of the effect of total yearly precipitation and mean yearly temperature on the proportion of annuals among all species in ecoregions with sufficient data for the four most annual-rich families. (**A**) Asteraceae, N = 510. (**B**) Brassicaceae N = 265. (**C**) Fabaceae N = 527. (**D**) Poaceae N = 517.

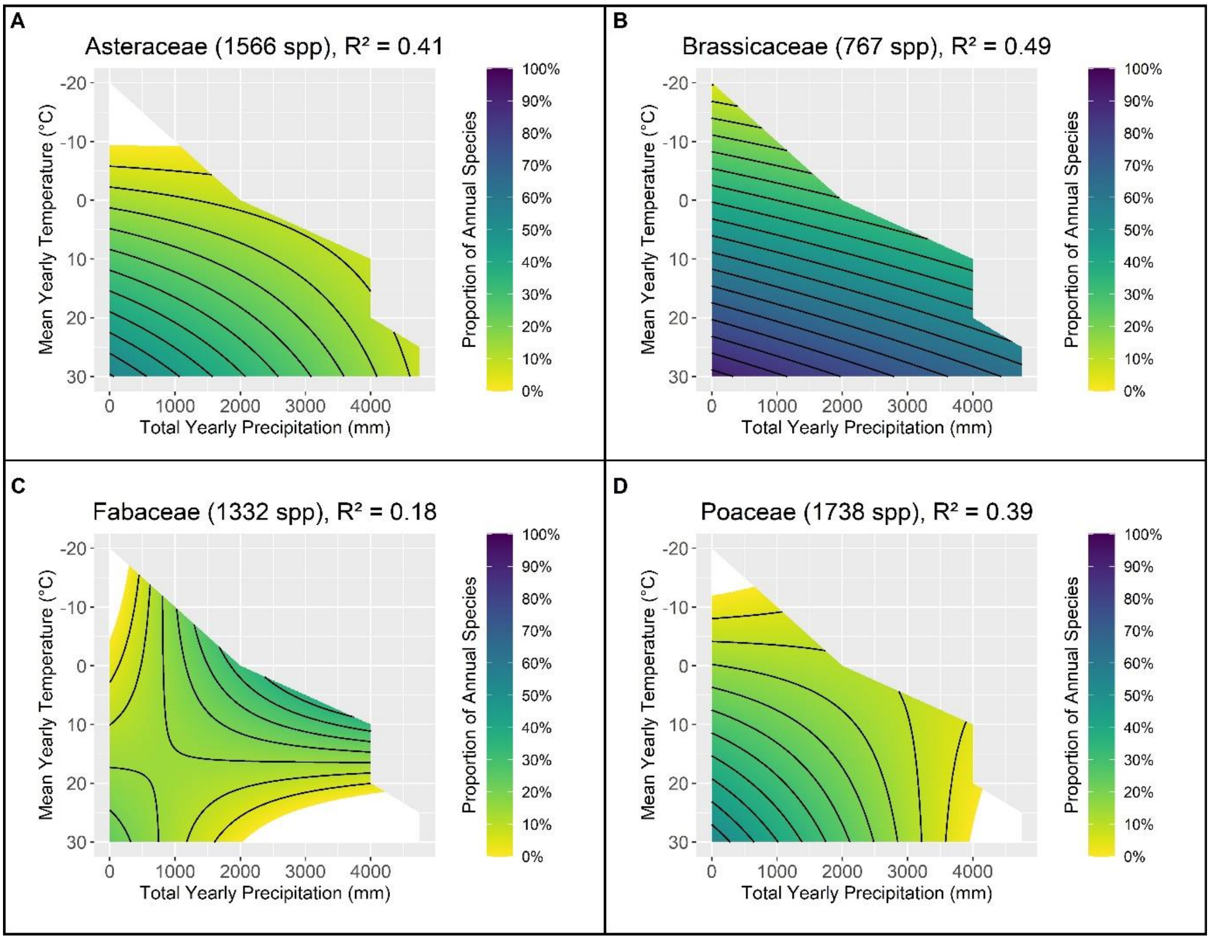


**Fig. S9.**
The effects of mean yearly precipitation and temperature on the proportion of annuals among all species in the four most annual-rich families (predictions of the regression model). (**A**) Asteraceae, N = 510. (**B**) Brassicaceae N = 265. (**C**) Fabaceae N = 527. (**D**) Poaceae N = 517. Contour lines are drawn every 5% for all figures.

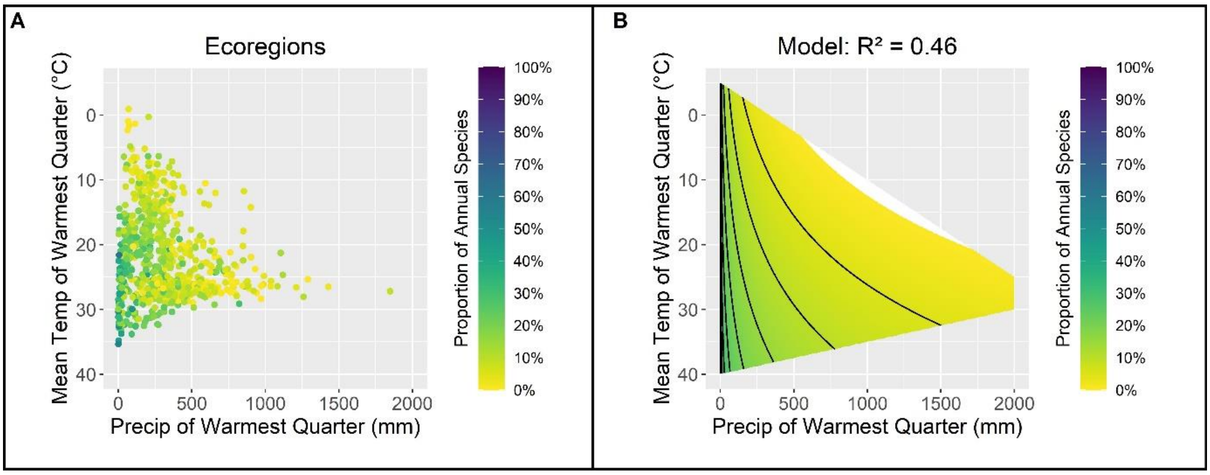


**Fig. S10.**
The effects of the total precipitation and mean temperature of the warmest quarter on the proportion of annuals among all species. (**A**) The annuals proportion in each ecoregion. (**B**) The predictions of a regression model of the annuals proportion as a function of precipitation and mean temperature of the warmest quarter with contour lines every 5%. N=726.

**Table S1.**
The results of a linear regression model assessing the relationship between the relative proportion of annual species among herbaceous plants and mean yearly temperature (BioClim_1) and total yearly precipitation (BioClim_12) in ecoregions with sufficient data.

| **Predictors** | **Estimate** | **Std Error** | **Statistics** | **P-value** |
|---|---|---|---|---|
| (Intercept) | 0.180637 | 0.011843 | 15.25232 | 2.19E-45 |
| BioClim_1 | 0.011238 | 0.000633 | 17.76618 | 2.79E-58 |
| BioClim_12 | -7.07E-05 | 1.56E-05 | -4.52991 | 6.97E-06 |
| BioClim_1:BioClim_12 | -2.13E-06 | 6.75E-07 | -3.15431 | 0.00168 |
| **Observations** | 683 | | | |
| **R-squared** | 0.4824 | | | |

**Table S2.**
The results of a linear regression model assessing the relationship between the relative proportion of annuals among herbaceous species in the Asteraceae family (1566 spp.) and mean yearly temperature (BioClim_1) and total yearly precipitation (BioClim_12) in ecoregions with sufficient data.

| Predictors | Estimate | Std Error | Statistics | P-value |
|---|---|---|---|---|
| (Intercept) | 0.186972 | 0.01648 | 11.34529 | 1.77E-26 |
| BioClim_1 | 0.020114 | 0.000971 | 20.71111 | 7.61E-68 |
| BioClim_12 | -8.62E-05 | 2.19E-05 | -3.94222 | 9.33E-05 |
| BioClim_1:BioClim_12 | 3.82E-07 | 1.05E-06 | 0.364298 | 0.715803 |
| **Observations** | 465 | | | |
| **R-squared** | 0.6833 | | | |

**Table S3.**
The results of a linear regression model assessing the relationship between the relative proportion of annuals among herbaceous species in the Brassicaceae family (767 spp.) and mean yearly temperature (BioClim_1) and total yearly precipitation (BioClim_12) in ecoregions with sufficient data.

| Predictors | Estimate | Std Error | Statistics | P-value |
|---|---|---|---|---|
| (Intercept) | 0.412271 | 0.022587 | 18.2526 | 2.33E-48 |
| BioClim_1 | 0.019554 | 0.001606 | 12.17787 | 2.95E-27 |
| BioClim_12 | -5.86E-05 | 3.70E-05 | -1.58383 | 0.114459 |
| BioClim_1:BioClim_12 | -1.52E-06 | 2.70E-06 | -0.56501 | 0.572559 |
| **Observations** | 262 | | | |
| **R-squared** | 0.5502 | | | |

**Table S4.**
The results of a linear regression model assessing the relationship between the relative proportion of annuals among herbaceous species in the Fabaceae family (1332 spp.) and mean yearly temperature (BioClim_1) and total yearly precipitation (BioClim_12) in ecoregions with sufficient data.

| Predictors | Estimate | Std Error | Statistics | P-value |
|---|---|---|---|---|
| (Intercept) | 0.045123 | 0.028561 | 1.579865 | 0.114974 |
| BioClim_1 | 0.021967 | 0.001694 | 12.96391 | 4.71E-32 |
| BioClim_12 | 0.000181 | 3.82E-05 | 4.739899 | 3.03E-06 |
| BioClim_1:BioClim_12 | -1.50E-05 | 1.85E-06 | -8.12816 | 6.29E-15 |
| **Observations** | 382 | | | |
| **R-squared** | 0.3175 | | | |

**Table S5.**
The results of a linear regression model assessing the relationship between the relative proportion of annuals among herbaceous species in the Poaceae family (1738 spp.) and mean yearly temperature (BioClim_1) and total yearly precipitation (BioClim_12) in ecoregions with sufficient data.

| Predictors | Estimate | Std Error | Statistics | P-value |
|---|---|---|---|---|
| (Intercept) | 0.154678 | 0.015717 | 9.841408 | 4.89E-21 |
| BioClim_1 | 0.013027 | 0.000883 | 14.75871 | 2.72E-41 |
| BioClim_12 | -1.92E-05 | 2.06E-05 | -0.93498 | 0.350243 |
| BioClim_1:BioClim_12 | -3.80E-06 | 9.36E-07 | -4.05596 | 5.78E-05 |
| **Observations** | 513 | | | |
| **R-squared** | 0.3868 | | | |

**Table S6.**

The results of a linear regression model assessing the relationship between the relative proportion of annuals among herbaceous species in ecoregions with sufficient data and mean temperature during the warmest quarter (BioClim_10) and the log transformation of the total precipitation during the warmest quarter (BioClim_18) in ecoregions with sufficient data.

| **Predictors** | **Estimate** | **Std Error** | **Statistics** | **P-value** |
|---|---|---|---|---|
| (Intercept) | 0.270432 | 0.076207 | 3.548666 | 0.000414 |
| BioClim_10 | 0.013782 | 0.002909 | 4.737117 | 2.64E-06 |
| log10(BioClim_18 + 1) | -0.12593 | 0.034049 | -3.6985 | 0.000234 |
| BioClim_10:log10(BioClim_18 + 1) | -0.00099 | 0.001307 | -0.76074 | 0.447078 |
| **Observations** | 683 | | | |
| **R-squared** | 0.5482 | | | |

**Table S7.**
The results of a linear regression model assessing the relationship between the relative proportion of annuals among herbaceous species and interannual precipitation variation (Inter_CoV) in ecoregions with sufficient data.

| Predictors | Estimate | Std Error | Statistics | P-value |
|---|---|---|---|---|
| (Intercept) | 0.058074 | 0.012555 | 4.625491 | 4.47E-06 |
| Inter_CoV | 0.010001 | 0.000677 | 14.78138 | 4.46E-43 |
| **Observations** | 682 | | | |
| **R-squared** | 0.2432 | | | |

**Table S8.**
The results of a linear regression model assessing the relationship between the relative proportion of annuals among herbaceous species and mean yearly temperature (BioClim_1), total yearly precipitation (BioClim_12), and interannual precipitation variation (Inter_CoV) in ecoregions with sufficient data.

| Predictors | Estimate | Std Error | Statistics | P-value |
|---|---|---|---|---|
| (Intercept) | 0.077734 | 0.036354 | 2.138261 | 0.032855 |
| BioClim_1 | 0.011485 | 0.001879 | 6.113557 | 1.65E-09 |
| BioClim_12 | -4.73E-05 | 4.60E-05 | -1.02823 | 0.304209 |
| Inter_CoV | 0.005691 | 0.002234 | 2.547546 | 0.011069 |
| BioClim_1:BioClim_12 | -1.23E-06 | 2.05E-06 | -0.59924 | 0.549212 |
| BioClim_1:Inter_CoV | -7.27E-05 | 0.000109 | -0.66793 | 0.504409 |
| BioClim_12:Inter_CoV | 4.11E-07 | 3.37E-06 | 0.122126 | 0.902836 |
| BioClim_1:BioClim_12:Inter_CoV | -9.60E-08 | 1.48E-07 | -0.64794 | 0.517241 |
| **Observations** | 682 | | | |
| **R-squared** | 0.5053 | | | |

**Table S9.**
The results of a linear regression model assessing the relationship between the relative proportion of annuals among herbaceous species and mean temperature during the warmest quarter (BioClim_10), the log transformation of the total precipitation during the warmest quarter (BioClim_18), and interannual precipitation variation (Inter_CoV) in ecoregions with sufficient data.

| Predictors | Estimate | Std Error | Statistics | P-value |
|---|---|---|---|---|
| (Intercept) | -0.10295 | 0.176862 | -0.58208 | 0.560708 |
| BioClim_10 | 0.030317 | 0.007331 | 4.135556 | 3.99E-05 |
| log10(BioClim_18 + 1) | 0.045801 | 0.07655 | 0.598312 | 0.549833 |
| Inter_CoV | 0.014033 | 0.007138 | 1.965808 | 0.049731 |
| BioClim_10:log10(BioClim_18 + 1) | -0.00979 | 0.003179 | -3.08014 | 0.002153 |
| BioClim_10:Inter_CoV | -0.0007 | 0.000294 | -2.37138 | 0.018002 |
| log10(BioClim_18 + 1):Inter_CoV | -0.00649 | 0.003406 | -1.90522 | 0.057177 |
| BioClim_10:log10(BioClim_18 + 1):Inter_CoV | 0.000393 | 0.000139 | 2.816811 | 0.004992 |
| **Observations** | 682 | | | |
| **R-squared** | 0.5832 | | | |

**Table S10.**
The results of a linear regression model assessing the relationship between the relative proportion of annuals among herbaceous species and human footprint (HumanF) in ecoregions with sufficient data.

| Predictors | Estimate | Std Error | Statistics | P-value |
|---|---|---|---|---|
| (Intercept) | 0.174181 | 0.011091 | 15.7051 | 1.16E-47 |
| HumanF | 0.002714 | 0.000476 | 5.706035 | 1.73E-08 |
| **Observations** | 683 | | | |
| **R-squared** | 0.0456 | | | |

**Table S11.**
The results of a linear regression model assessing the relationship between the relative proportion of annuals among herbaceous species and mean yearly temperature (BioClim_1), total yearly precipitation (BioClim_12), and human footprint (HumanF) in ecoregions with sufficient data.

| Predictors | Estimate | Std Error | Statistics | P-value |
|---|---|---|---|---|
| (Intercept) | 0.155262 | 0.017988 | 8.631379 | 4.34E-17 |
| BioClim_1 | 0.012907 | 0.000965 | 13.36809 | 2.50E-36 |
| BioClim_12 | -8.08E-05 | 2.66E-05 | -3.03353 | 0.00251 |
| HumanF | 0.003448 | 0.001307 | 2.638072 | 0.00853 |
| BioClim_1:BioClim_12 | -2.59E-06 | 1.13E-06 | -2.29337 | 0.022133 |
| BioClim_1:HumanF | -0.0002 | 6.42E-05 | -3.18806 | 0.001498 |
| BioClim_12:HumanF | -7.26E-07 | 1.53E-06 | -0.47472 | 0.635143 |
| BioClim_1:BioClim_12:HumanF | 8.52E-08 | 6.44E-08 | 1.321897 | 0.18665 |
| **Observations** | 683 | | | |
| **R-squared** | 0.5004 | | | |

**Table S12.**
The results of a linear regression model assessing the relationship between the relative proportion of annuals among herbaceous species and mean temperature during the warmest quarter (BioClim_10), the log transformation of the total precipitation during the warmest quarter (BioClim_18), and human footprint (HumanF) in ecoregions with sufficient data.

| Predictors | Estimate | Std Error | Statistics | P-value |
|---|---|---|---|---|
| (Intercept) | 0.069872 | 0.149197 | 0.468319 | 0.639708 |
| BioClim_10 | 0.02225 | 0.005577 | 3.989825 | 7.34E-05 |
| log10(BioClim_18 + 1) | -0.05578 | 0.067294 | -0.82883 | 0.407492 |
| HumanF | 0.015269 | 0.007958 | 1.918651 | 0.05545 |
| BioClim_10:log10(BioClim_18 + 1) | -0.00443 | 0.002538 | -1.74611 | 0.081247 |
| BioClim_10:HumanF | -0.00062 | 0.000303 | -2.03609 | 0.042132 |
| log10(BioClim_18 + 1):HumanF | -0.00487 | 0.003509 | -1.38838 | 0.165478 |
| BioClim_10:log10(BioClim_18 + 1):HumanF | 0.000216 | 0.000134 | 1.617571 | 0.106222 |
| **Observations** | 683 | | | |
| **R-squared** | 0.5659 | | | |

**Table S13.**
The results of a linear regression model assessing the relationship between the relative proportion of annuals among herbaceous species and mean yearly temperature (BioClim_1), total yearly precipitation (BioClim_12), interannual precipitation variation (Inter_CoV), and human footprint (HumanF) in ecoregions with sufficient data.

| Predictors | Estimate | Std Error | Statistics | P-value |
|---|---|---|---|---|
| (Intercept) | 0.231736 | 0.06039 | 3.837322 | 0.000136 |
| BioClim_1 | 0.004947 | 0.003159 | 1.566119 | 0.117796 |
| BioClim_12 | -0.0002 | 8.19E-05 | -2.4126 | 0.016108 |
| Inter_CoV | -0.0064 | 0.003979 | -1.60929 | 0.108026 |
| HumanF | -0.01134 | 0.004466 | -2.54007 | 0.011309 |
| BioClim_1:BioClim_12 | 3.91E-06 | 3.62E-06 | 1.080932 | 0.280119 |
| BioClim_1:Inter_CoV | 0.000442 | 0.00019 | 2.322801 | 0.02049 |
| BioClim_12:Inter_CoV | 9.79E-06 | 6.32E-06 | 1.54971 | 0.121686 |
| BioClim_1:HumanF | 0.000433 | 0.000207 | 2.089613 | 0.037031 |
| BioClim_12:HumanF | 1.20E-05 | 5.10E-06 | 2.353931 | 0.018866 |
| Inter_CoV:HumanF | 0.000958 | 0.000278 | 3.446795 | 0.000603 |
| BioClim_1:BioClim_12:Inter_CoV | -4.64E-07 | 2.78E-07 | -1.67188 | 0.095017 |
| BioClim_1:BioClim_12:HumanF | -4.15E-07 | 2.16E-07 | -1.91938 | 0.055363 |
| BioClim_1:Inter_CoV:HumanF | -3.96E-05 | 1.26E-05 | -3.14805 | 0.001717 |
| BioClim_12:Inter_CoV:HumanF | -8.51E-07 | 3.57E-07 | -2.38167 | 0.017514 |
| BioClim_1:BioClim_12:Inter_CoV:HumanF | 3.31E-08 | 1.52E-08 | 2.179352 | 0.029655 |
| **Observations** | 682 | | | |
| **R-squared** | 0.532 | | | |

**Table S14.**

The results of a linear regression model assessing the relationship between the relative proportion of annuals among herbaceous species and mean temperature during the warmest quarter (BioClim_10), the log transformation of the total precipitation during the warmest quarter (BioClim_18), interannual precipitation variation (Inter_CoV), and human footprint (HumanF) in ecoregions with sufficient data.

| Predictors | Estimate | Std Error | Statistics | P-value |
|---|---|---|---|---|
| (Intercept) | -0.09761 | 0.339092 | -0.28785 | 0.773553 |
| BioClim_10 | 0.026529 | 0.013463 | 1.970523 | 0.049192 |
| log10(BioClim_18 + 1) | 0.031909 | 0.146687 | 0.217533 | 0.82786 |
| Inter_CoV | 0.00083 | 0.016006 | 0.051842 | 0.95867 |
| HumanF | 0.00712 | 0.019725 | 0.360951 | 0.718251 |
| BioClim_10:log10(BioClim_18 + 1) | -0.00913 | 0.005843 | -1.56303 | 0.118519 |
| BioClim_10:Inter_CoV | -0.0001 | 0.000628 | -0.16351 | 0.870165 |
| log10(BioClim_18 + 1):Inter_CoV | -0.00037 | 0.007352 | -0.0505 | 0.959736 |
| BioClim_10:HumanF | -0.00014 | 0.00076 | -0.17958 | 0.857535 |
| log10(BioClim_18 + 1):HumanF | -0.00016 | 0.00855 | -0.01852 | 0.985233 |
| Inter_CoV:HumanF | 0.000612 | 0.000934 | 0.654874 | 0.512775 |
| BioClim_10:log10(BioClim_18 + 1):Inter_CoV | 0.000166 | 0.000289 | 0.57517 | 0.56537 |
| BioClim_10:log10(BioClim_18 + 1):HumanF | 1.34E-05 | 0.000329 | 0.040574 | 0.967648 |
| BioClim_10:Inter_CoV:HumanF | -2.58E-05 | 3.51E-05 | -0.73461 | 0.462834 |
| log10(BioClim_18 + 1):Inter_CoV:HumanF | -0.00038 | 0.000429 | -0.87931 | 0.379552 |
| BioClim_10:log10(BioClim_18 + 1):Inter_CoV:HumanF | 1.31E-05 | 1.62E-05 | 0.807203 | 4.20E-01 |
| **Observations** | 682 | | | |
| **R-squared** | 0.6071 | | | |

**Table S15.**
The results of a linear regression model assessing the relationship between the relative proportion of annuals among all plant species and mean yearly temperature (BioClim_1) and total yearly precipitation (BioClim_12) in ecoregions with sufficient data.

| Predictors | Estimate | Std Error | Statistics | P-value |
|---|---|---|---|---|
| (Intercept) | 0.1254 | 0.009221 | 13.59997 | 1.14E-37 |
| BioClim_1 | 0.005098 | 0.000486 | 10.49588 | 4.41E-24 |
| BioClim_12 | -2.11E-05 | 1.21E-05 | -1.74227 | 0.081888 |
| BioClim_1:BioClim_12 | -2.60E-06 | 5.18E-07 | -5.02685 | 6.29E-07 |
| **Observations** | 726 | | | |
| **R-squared** | 0.3416 | | | |

**Table S16.**

The results of a linear regression model assessing the relationship between the relative proportion of annuals among all plant species in the Asteraceae family (1566 spp.) and mean yearly temperature (BioClim_1) and total yearly precipitation (BioClim_12) in ecoregions with sufficient data.

| Predictors | Estimate | Std Error | Statistics | P-value |
|---|---|---|---|---|
| (Intercept) | 0.131708 | 0.016751 | 7.86276 | 2.28E-14 |
| BioClim_1 | 0.014138 | 0.000971 | 14.5655 | 2.19E-40 |
| BioClim_12 | -2.39E-05 | 2.16E-05 | -1.10368 | 0.270257 |
| BioClim_1:BioClim_12 | -2.51E-06 | 1.01E-06 | -2.4848 | 0.013285 |
| **Observations** | 510 | | | |
| **R-squared** | 0.4138 | | | |

**Table S17.**

The results of a linear regression model assessing the relationship between the relative proportion of annuals among all plant species in the Brassicaceae family (767 spp.) and mean yearly temperature (BioClim_1) and total yearly precipitation (BioClim_12) in ecoregions with sufficient data.

| **Predictors** | **Estimate** | **Std Error** | **Statistics** | **P-value** |
|---|---|---|---|---|
| (Intercept) | 0.394659 | 0.023134 | 17.05947 | 2.38E-44 |
| BioClim_1 | 0.017515 | 0.001633 | 10.72404 | 1.80E-22 |
| BioClim_12 | -4.49E-05 | 3.79E-05 | -1.18264 | 0.238029 |
| BioClim_1:BioClim_12 | -5.35E-07 | 2.76E-06 | -0.19372 | 0.846543 |
| **Observations** | 265 | | | |
| **R-squared** | 0.4945 | | | |

**Table S18.**
The results of a linear regression model assessing the relationship between the relative proportion of annuals among all plant species in the Fabaceae family (1332 spp.) and mean yearly temperature (BioClim_1) and total yearly precipitation (BioClim_12) in ecoregions with sufficient data.

| Predictors | Estimate | Std Error | Statistics | P-value |
|---|---|---|---|---|
| (Intercept) | 0.030548 | 0.018011 | 1.696047 | 0.090472 |
| BioClim_1 | 0.006903 | 0.000949 | 7.276429 | 1.26E-12 |
| BioClim_12 | 0.000146 | 2.32E-05 | 6.282645 | 7.01E-10 |
| BioClim_1:BioClim_12 | -8.76E-06 | 9.96E-07 | -8.79346 | 2.10E-17 |
| **Observations** | 527 | | | |
| **R-squared** | 0.176 | | | |

**Table S19.**
The results of a linear regression model assessing the relationship between the relative proportion of annuals among all plant species in the Poaceae family (1738 spp.) and mean yearly temperature (BioClim_1) and total yearly precipitation (BioClim_12) in ecoregions with sufficient data.

| Predictors | Estimate | Std Error | Statistics | P-value |
|---|---|---|---|---|
| (Intercept) | 0.152966 | 0.015328 | 9.979625 | 1.49E-21 |
| BioClim_1 | 0.012844 | 0.000862 | 14.89767 | 5.70E-42 |
| BioClim_12 | -2.13E-05 | 2.01E-05 | -1.0636 | 0.288008 |
| BioClim_1:BioClim_12 | -3.83E-06 | 9.14E-07 | -4.19289 | 3.24E-05 |
| **Observations** | 517 | | | |
| **R-squared** | 0.3911 | | | |

**Table S20.**
The results of a linear regression model assessing the relationship between the relative proportion of annuals among all plant species in ecoregions with sufficient data and mean temperature during the warmest quarter (BioClim_10) and the log transformation of the total precipitation during the warmest quarter (BioClim_18) in ecoregions with sufficient data.

| Predictors | Estimate | Std Error | Statistics | P-value |
|---|---|---|---|---|
| (Intercept) | 0.213172 | 0.05712 | 3.73198 | 0.000205 |
| BioClim_10 | 0.008619 | 0.002169 | 3.974695 | 7.76E-05 |
| log10(BioClim_18 + 1) | -0.07052 | 0.025523 | -2.76301 | 0.005873 |
| BioClim_10:log10(BioClim_18 + 1) | -0.00228 | 0.000974 | -2.34484 | 0.019305 |
| **Observations** | 726 | | | |
| **R-squared** | 0.4593 | | | |

**Table S21.**

The results of a linear regression model assessing the relationship between the relative proportion of annuals among all plant species and interannual precipitation variation (Inter_CoV) in ecoregions with sufficient data.

| Predictors | Estimate | Std Error | Statistics | P-value |
|---|---|---|---|---|
| (Intercept) | 0.027107 | 0.008529 | 3.178092 | 0.001546 |
| Inter_CoV | 0.005548 | 0.000452 | 12.27657 | 1.33E-31 |
| **Observations** | 725 | | | |
| **R-squared** | 0.1725 | | | |

**Table S22.**
The results of a linear regression model assessing the relationship between the relative proportion of annuals among all plant species and mean yearly temperature (BioClim_1), total yearly precipitation (BioClim_12), and interannual precipitation variation (Inter_CoV) in ecoregions with sufficient data.

| Predictors | Estimate | Std Error | Statistics | P-value |
|---|---|---|---|---|
| (Intercept) | 0.036436 | 0.02813 | 1.295273 | 0.195643 |
| BioClim_1 | 0.007012 | 0.001363 | 5.145224 | 3.45E-07 |
| BioClim_12 | -3.27E-05 | 3.43E-05 | -0.95203 | 0.341401 |
| Inter_CoV | 0.004859 | 0.0017 | 2.859025 | 0.004373 |
| BioClim_1:BioClim_12 | -1.12E-06 | 1.50E-06 | -0.75166 | 0.4525 |
| BioClim_1:Inter_CoV | -0.00014 | 7.85E-05 | -1.73947 | 0.082381 |
| BioClim_12:Inter_CoV | 2.74E-06 | 2.44E-06 | 1.123676 | 0.261527 |
| BioClim_1:BioClim_12:Inter_CoV | -1.59E-07 | 1.06E-07 | -1.4991 | 0.134289 |
| **Observations** | 725 | | | |
| **R-squared** | 0.371 | | | |

**Table S23.**
The results of a linear regression model assessing the relationship between the relative proportion of annuals among all plant species and mean temperature during the warmest quarter (BioClim_10), the log transformation of the total precipitation during the warmest quarter (BioClim_18), and interannual precipitation variation (Inter_CoV) in ecoregions with sufficient data.

| Predictors | Estimate | Std Error | Statistics | P-value |
|---|---|---|---|---|
| (Intercept) | -0.03287 | 0.128753 | -0.25532 | 0.798551 |
| BioClim_10 | 0.023532 | 0.005069 | 4.642038 | 4.10E-06 |
| log10(BioClim_18 + 1) | 0.036468 | 0.055866 | 0.652768 | 0.514115 |
| Inter_CoV | 0.008155 | 0.005135 | 1.588326 | 0.112653 |
| BioClim_10:log10(BioClim_18 + 1) | -0.00925 | 0.002212 | -4.18177 | 3.25E-05 |
| BioClim_10:Inter_CoV | -0.00056 | 0.000197 | -2.84618 | 0.004551 |
| log10(BioClim_18 + 1):Inter_CoV | -0.00354 | 0.002444 | -1.44954 | 0.147623 |
| BioClim_10:log10(BioClim_18 + 1):Inter_CoV | 0.000279 | 9.47E-05 | 2.940618 | 0.003381 |
| **Observations** | 725 | | | |
| **R-squared** | 0.4959 | | | |

**Table S24.**
The results of a linear regression model assessing the relationship between the relative proportion of annuals among all plant species and human footprint (HumanF) in ecoregions with sufficient data.

| Predictors | Estimate | Std Error | Statistics | P-value |
|---|---|---|---|---|
| (Intercept) | 0.105067 | 0.007747 | 13.56224 | 1.68E-37 |
| HumanF | 0.000862 | 0.000329 | 2.617391 | 0.009045 |
| **Observations** | 726 | | | |
| **R-squared** | 0.0094 | | | |

**Table S25.**
The results of a linear regression model assessing the relationship between the relative proportion of annuals among all plant species and mean yearly temperature (BioClim_1), total yearly precipitation (BioClim_12), and human footprint (HumanF) in ecoregions with sufficient data.

| Predictors | Estimate | Std Error | Statistics | P-value |
|---|---|---|---|---|
| (Intercept) | 0.087571 | 0.013755 | 6.366423 | 3.45E-10 |
| BioClim_1 | 0.007491 | 0.000732 | 10.22961 | 5.07E-23 |
| BioClim_12 | -3.59E-05 | 2.04E-05 | -1.76416 | 0.078131 |
| HumanF | 0.004999 | 0.000991 | 5.045102 | 5.75E-07 |
| BioClim_1:BioClim_12 | -2.51E-06 | 8.59E-07 | -2.9252 | 0.003551 |
| BioClim_1:HumanF | -0.00028 | 4.79E-05 | -5.89369 | 5.81E-09 |
| BioClim_12:HumanF | -1.04E-06 | 1.16E-06 | -0.90191 | 0.367405 |
| BioClim_1:BioClim_12:HumanF | 8.39E-08 | 4.85E-08 | 1.730612 | 0.083951 |
| **Observations** | 726 | | | |
| **R-squared** | 0.3977 | | | |

**Table S26.**
The results of a linear regression model assessing the relationship between the relative proportion of annuals among all plant species and mean temperature during the warmest quarter (BioClim_10), the log transformation of the total precipitation during the warmest quarter (BioClim_18), and human footprint (HumanF).

| Predictors | Estimate | Std Error | Statistics | P-value |
|---|---|---|---|---|
| (Intercept) | 0.01041 | 0.111548 | 0.093319 | 0.925676 |
| BioClim_10 | 0.015733 | 0.004136 | 3.803779 | 0.000155 |
| log10(BioClim_18 + 1) | -0.00768 | 0.050329 | -0.15263 | 0.878735 |
| HumanF | 0.015163 | 0.005835 | 2.598751 | 0.009548 |
| BioClim_10:log10(BioClim_18 + 1) | -0.00461 | 0.001883 | -2.44776 | 0.014613 |
| BioClim_10:HumanF | -0.00055 | 0.00022 | -2.47181 | 0.013674 |
| log10(BioClim_18 + 1):HumanF | -0.00435 | 0.002577 | -1.68673 | 0.09209 |
| BioClim_10:log10(BioClim_18 + 1):HumanF | 0.000159 | 9.77E-05 | 1.626196 | 0.104347 |
| **Observations** | 726 | | | |
| **R-squared** | 0.4946 | | | |

**Table S27.**
The results of a linear regression model assessing the relationship between the relative proportion of annuals among all plant species and mean yearly temperature (BioClim_1), total yearly precipitation (BioClim_12), interannual precipitation variation (Inter_CoV), and human footprint (HumanF) in ecoregions with sufficient data.

| Predictors | Estimate | Std Error | Statistics | P-value |
|---|---|---|---|---|
| (Intercept) | 0.207773 | 0.045498 | 4.566631 | 5.84E-06 |
| BioClim_1 | -0.0005 | 0.002296 | -0.2184 | 0.827182 |
| BioClim_12 | -0.00019 | 6.14E-05 | -3.10947 | 0.001949 |
| Inter_CoV | -0.00905 | 0.002971 | -3.04705 | 0.002397 |
| HumanF | -0.01096 | 0.003254 | -3.36894 | 0.000795 |
| BioClim_1:BioClim_12 | 4.92E-06 | 2.66E-06 | 1.849078 | 0.064863 |
| BioClim_1:Inter_CoV | 0.000488 | 0.000138 | 3.539681 | 0.000427 |
| BioClim_12:Inter_CoV | 1.23E-05 | 4.72E-06 | 2.608674 | 0.009281 |
| BioClim_1:HumanF | 0.000413 | 0.000141 | 2.922374 | 0.003584 |
| BioClim_12:HumanF | 1.16E-05 | 3.59E-06 | 3.242638 | 0.00124 |
| Inter_CoV:HumanF | 0.001021 | 0.000199 | 5.124802 | 3.84E-07 |
| BioClim_1:BioClim_12:Inter_CoV | -5.47E-07 | 2.04E-07 | -2.68373 | 0.007451 |
| BioClim_1:BioClim_12:HumanF | -4.30E-07 | 1.49E-07 | -2.88948 | 0.003977 |
| BioClim_1:Inter_CoV:HumanF | -4.34E-05 | 8.51E-06 | -5.10478 | 4.26E-07 |
| BioClim_12:Inter_CoV:HumanF | -8.43E-07 | 2.46E-07 | -3.43309 | 0.000631 |
| BioClim_1:BioClim_12:Inter_CoV:HumanF | 3.40E-08 | 1.03E-08 | 3.308154 | 0.000987 |
| **Observations** | 725 | | | |
| **R-squared** | 0.4363 | | | |

**Table S28.**
The results of a linear regression model assessing the relationship between the relative proportion of annuals among all plant species and mean temperature during the warmest quarter (BioClim_10), the log transformation of the total precipitation during the warmest quarter (BioClim_18), interannual precipitation variation (Inter_CoV), and human footprint (HumanF) in ecoregions with sufficient data.

| Predictors | Estimate | Std Error | Statistics | P-value |
|---|---|---|---|---|
| (Intercept) | 0.04513 | 0.244929 | 0.184259 | 0.853863 |
| BioClim_10 | 0.019474 | 0.009155 | 2.127066 | 0.033759 |
| log10(BioClim_18 + 1) | -0.00652 | 0.105893 | -0.06155 | 0.950942 |
| Inter_CoV | -0.00545 | 0.011278 | -0.483 | 0.629244 |
| HumanF | 0.004122 | 0.013843 | 0.297743 | 0.765987 |
| BioClim_10:log10(BioClim_18 + 1) | -0.00825 | 0.003983 | -2.07071 | 0.038747 |
| BioClim_10:Inter_CoV | -7.29E-05 | 0.00041 | -0.17769 | 0.859015 |
| log10(BioClim_18 + 1):Inter_CoV | 0.001758 | 0.005187 | 0.338956 | 0.734743 |
| BioClim_10:HumanF | -0.00016 | 0.000511 | -0.30618 | 0.759558 |
| log10(BioClim_18 + 1):HumanF | 0.000916 | 0.006041 | 0.151631 | 0.879521 |
| Inter_CoV:HumanF | 0.000606 | 0.00065 | 0.931993 | 0.351657 |
| BioClim_10:log10(BioClim_18 + 1):Inter_CoV | 0.000136 | 0.00019 | 0.714869 | 0.474925 |
| BioClim_10:log10(BioClim_18 + 1):HumanF | 1.54E-05 | 0.000225 | 0.068304 | 0.945563 |
| BioClim_10:Inter_CoV:HumanF | -1.94E-05 | 2.33E-05 | -0.83523 | 0.403869 |
| log10(BioClim_18 + 1):Inter_CoV:HumanF | -0.00031 | 0.000298 | -1.05026 | 0.293957 |
| BioClim_10:log10(BioClim_18 + 1):Inter_CoV:HumanF | 8.03E-06 | 1.09E-05 | 0.738153 | 0.460666 |
| **Observations** | 725 | | | |
| **R-squared** | 0.5406 | | | |

**Table S29.**
The relative proportion of annuals among all plant species and among herbaceous species in each of the biomes defined by (*37*). See methods for biome classification.

| Biome | Annual Frequency | Annual Herb Frequency |
|---|---|---|
| Desert | 14% | 25% |
| Woodland / Grassland | 9% | 19% |
| Tropical Dry Forest | 3% | 14% |
| Tropical Wet Forest | 3% | 14% |
| Temperate Forest | 9% | 16% |
| Temperate Wet Forest | 7% | 14% |
| Boreal Forest | 11% | 14% |
| Tundra | 11% | 13% |

**Table S30.**
The previous estimates obtained from (*30*), the location of the estimates' original study, our corresponding biome based on the original study location, and the calculations used to determine the proportion of annuals among herbs from the available data. For information on the origin of these estimates see Table S33.

| **Initial Biome Nomenclature** (Begon & Townsend, 2020) | **Original Study Location** | **Corresponding Biome Nomenclature** | **Annuals among All Species** | **Annuals among Herbaceous Species** |
|---|---|---|---|---|
| Global | | Global | 13% | $\frac{13}{3+27+3+1+13} \approx 27.65\%$ |
| Arctic | Baffin's Island | Tundra | 2% | $\frac{2}{0+51+13+3+2} \approx 2.89\%$ |
| Desert | Death Valley | Desert | 42% | $\frac{42}{0+18+2+5+42} \approx 62.68\%$ |
| Tropical | Seychelles | Wet tropical forest | 16% | $\frac{16}{3+12+3+2+16} \approx 44.44\%$ |
| Temperate | Denmark | Temperate Forest | 18% | $\frac{18}{0+50+11+11+18} = 20\%$ |
| Mediterranean | The Camargue (mouth of the Rhone) | Woodland/ Grassland | 39% | $\frac{39}{0+23+11+39} \approx 53.42\%$ |
| The proportion of annuals among Herbaceous Species is determined from values taken from the original studies and is calculated by: $\frac{Therophyte}{Epiphyte + Hemicryptophyte + Geophyte + Hydrophyte + Therophyte}$ | | | | |

**Table S31.**
A comparison of previous estimates, obtained from (*31*), for the proportion of annuals among all species and among herbs to our revised estimates. Greyed cells have no initial biome estimate. Alternative previous estimates from (*30*) are available in Table 1. Note, the biome nomenclature used for the previous estimates differs from ours and so the location of the original study was used to determine our corresponding biome. Additional information can be found in Table S32.

| | **Annuals among All Species** | | **Annuals among Herbaceous Species** | |
|---|---|---|---|---|
| Region | Previous Estimate | Revised Estimate | Previous Estimate | Revised Estimate |
| **Global** | **13%** | **6%** | **29%** | **13%** |
| Desert | 73% or 27% | 14% | 76%, 66% | 25% |
| Tundra | 2% | 11% | 3% | 13% |
| Woodland / Grassland | 4%, 6%, 14% | 9% | 11%, 13%, 16% | 19% |
| Boreal forest | | 11% | | 14% |
| Tropical Dry Forest | 0% | 3% | 0% | 14% |
| Tropical Wet Forest | 10% | 3% | 59% | 14% |
| Temperate Forest | 7%, 2% | 9% | 10%, 3% | 16% |
| Temperate Wet Forest | | 7% | | 14% |

**Table S32.**

The previous estimates obtained from (*31*), the location of the estimates' original study, our corresponding biome based on the original study location, and the calculations used to determine the proportion of annuals among herbs from the available data. For information on the origin of these estimates see Table S34.

| Initial Biome Nomenclature (Begon *et al.*, 2020) | Original Study Location | Corresponding Biome Nomenclature | Annuals among All Species | Annuals among Herbaceous Species |
|---|---|---|---|---|
| Global | | Global | 13% | $\frac{13}{26+6+13} \approx 28.88\%$ |
| Tropical Rainforest | Queensland | Tropical Dry Forest | 0% | 0% |
| Subtropical forest | Matheran, India | Tropical Wet Forest | 10% | $\frac{10}{2+5+10} \approx 58.82\%$ |
| Warm-temperate Forest | Mediterranean live-oak forest, 0-500 m | Woodland /Grassland | 4% | $\frac{4}{24+9+4} \approx 10.81\%$ |
| Cold-temperate forest | Central Siskiyou Mtns | Temperate Forest | 7% | $\frac{7}{54+12+7} \approx 9.58\%$ |
| Tundra | Spitzbergen | Tundra | 2% | $\frac{2}{60+15+2} \approx 2.59\%$ |
| Mid-temperate mesophytic forest | Siskiyou Mtns South Fork and Beaver Creek | Temperate Forest | 2% | $\frac{2}{33+23+2} \approx 3.44\%$ |
| Oak Woodland | Santa Catalina Mountains, Arizona | Woodland /Grassland | 6% | $\frac{6}{36+5+6} \approx 12.76\%$ |
| Dry Grassland | Pamir Mts. Steppe | Woodland /Grassland | 14% | $\frac{14}{63+10+14} \approx 16.09\%$ |
| Semi-desert | Oudjda semi-desert | Desert | 27% | $\frac{27}{14+0+27} \approx 65.85\%$ |
| Desert | Oudjda desert | Desert | 73% | $\frac{73}{17+6+73} \approx 76.04\%$ |

The proportion of annuals among Herbaceous Species is determined from values taken from the textbook (original study estimates were not accessible) and is calculated by:

$$\frac{Therophyte}{Hemicryptophyte + Cryptophyte + Therophyte}$$

**Table S33.**
The original studies of global and biome-level annuals proportion estimates found in (*29*, *30*), their location and sample size. Note that origin for the global annual frequency estimate is the same as in Table S34.

| Biome | Proportion | Original Study | Location | Species |
|---|---|---|---|---|
| Global | 13% | Raunkiær, C. (1918). Über das biologische Normalspektrum. Andr. Fred. Høst & søn, Bianco Lunos bogtrykkeri. | Worldwide | 400 |
| Tropical | 16% | Raunkiær, C. (1918). Über das biologische Normalspektrum. Andr. Fred. Høst & søn, Bianco Lunos bogtrykkeri. | Seychelles | 258 |
| Desert | 42% | Raunkiær, C. (1918). Über das biologische Normalspektrum. Andr. Fred. Høst & søn, Bianco Lunos bogtrykkeri. | Death Valley | 294 |
| Mediterranean | 39% | Raunkiær, C. (1918). Über das biologische Normalspektrum. Andr. Fred. Høst & søn, Bianco Lunos bogtrykkeri. | The Camargue (mouth of the Rhone) | 233 |
| Temperate | 18% | Raunkiær, C. (1918). Über das biologische Normalspektrum. Andr. Fred. Høst & søn, Bianco Lunos bogtrykkeri. | Denmark | 1084 |
| Arctic | 2% | Raunkiær, C. (1918). Über das biologische Normalspektrum. Andr. Fred. Høst & søn, Bianco Lunos bogtrykkeri. | Baffin’s Land | 129 |

**Table S34.**
The original studies of global and biome-level annuals proportion estimates found in (*28*, *31*), their location and sample size. Note that origin for the global annual frequency estimate is the same as in Table S33.

| Biome | Proportion | Original Study | Location | Species |
|---|---|---|---|---|
| Global | 13% | Raunkiær, C. (1918). Über das biologische Normalspektrum. Andr. Fred. Høst & søn, Bianco Lunos bogtrykkeri. | Worldwide | 400 |
| Tropical Rainforest | 0% | Cromer, D. A. N., & Pryor, L. D. (1942). A contribution to rain-forest ecology. In Proceedings of the Linnean Society (Vol. 67, pp. 249-268). | Queensland, Australia | 141 |
| Subtropical Forest | 10% | Bharucha, F. R., & Ferreira, D. B. (1941). The biological spectrum of the Madras flora. Journal of University of Bombay, 9, 93-100. | Matheran, India | 361 |
| Warm-temperate Forest | 4% | Braun-Blanquet, J., (1936). La Chênaie d'Yeuse méditerranéenne (Quercion ilicis): monographie phytosociologique. Sta. Internatl. Dr Géobot Méditer. Et Alp. Montpellier Commun. **45**: 1-147 | Mediterranean live-oak forest, 0-500 m | |
| Cold-temperate Forest | 7% | Whittaker, R. H. (1960). Vegetation of the Siskiyou Mountains, Oregon and California. Ecological Monographs, 30(3), 279–338. | Central Siskiyou Mtns., by elevation belts on diorite: 1920-2140 m | 72 |
| Tundra | 2% | Raunkiær, C. (1918). Über das biologische Normalspektrum. Andr. Fred. Høst & søn, Bianco Lunos bogtrykkeri. | Spitzbergen | 110 |
| Mid-temperate Forest | 2% | Whittaker, R. H. (1960). Vegetation of the Siskiyou Mountains, Oregon and California. Ecological Monographs, 30(3), 279–338. | Mixed Evergreen Forest in the West (South Fork and Beaver Creek) | 160 |
| Oak Woodland | 6% | Whittaker, R. H., & Niering, W. A. (1965). Vegetation of the Santa Catalina Mountains, Arizona: a gradient analysis of the south slope. Ecology, 46(4), 429-452. | Santa Catalina Mountains, Arizona | 100 |
| Dry Grassland | 14% | Paulsen, O. (1915). Some remarks on the desert vegetation of America. The Plant World, 18(6), 155-161. | Pamir Mts. Steppe | 514 |
| Semi-desert | 27% | Braun-Blanquet, J., & Maire, R. C. J. E. (1924). Etudes sur la végétation et la flore marocaines. | Oudjda semi-desert | 32 |
| Desert | 73% | Braun-Blanquet, J., & Maire, R. C. J. E. (1924). Etudes sur la végétation et la flore marocaines. | Oudjda desert | 49 |

**Table S35.**
The list of databases used in developing our growth-form database. The access date reflects the date the respective data was downloaded.

| Database Name | Access | Citation |
|---|---|---|
| BIEN | 01 June 2021 | B. S. Maitner, B. Boyle, N. Casler, R. Condit, J. Donoghue II, S. M. Durán, D. Guaderrama, C. E. Hinchliff, P. M. Jørgensen, N. J. B. Kraft, B. McGill, C. Merow, N. Morueta-Holme, R. K. Peet, B. Sandel, M. Schildhauer, S. A. Smith, J.-C. Svenning, B. Thiers, C. Violle, S. Wiser, B. J. Enquist, The bien r package: A tool to access the Botanical Information and Ecology Network (BIEN) database. *Methods Ecol. Evol.* **9**, 373–379 (2018). |
| BROT 2.0 | 23 May 2021 | Ç. Tavşanoğlu, J. G. Pausas, A functional trait database for Mediterranean Basin plants. *Sci. Data*. **5**, 180135 (2018). |
| EOL | 24 May 2021 | C. S. Parr, M. N. Wilson, M. P. Leary, K. S. Schulz, M. K. Lans, M. L. Walley, J. A. Hammock, M. A. Goddard, M. J. Rice, M. M. Studer, The encyclopedia of life v2: providing global access to knowledge about life on earth. *Biodivers. data J.* (2014). |
| Engemann *et al.*, 2016 | 03 June 2021 | K. Engemann, B. Sandel, B. Boyle, B. J. Enquist, P. M. Jørgensen, J. Kattge, B. J. McGill, N. Morueta-Holme, R. K. Peet, N. J. Spencer, C. Violle, S. K. Wiser, J. C. Svenning, A plant growth form dataset for the New World. *Ecology*. **97**, 3243 (2016). |
| Kew Gardens WCSP | 20 July 2021 | WCSP (2021). 'World Checklist of Selected Plant Families. Facilitated by the Royal Botanic Gardens, Kew. Published on the Internet; http://apps.kew.org/wcsp/ Retrieved July 20, 2021.'<br>**direct correspondence* |
| LEDA | 01 June 2021 | M. Kleyer, R. M. Bekker, I. C. Knevel, J. P. Bakker, K. Thompson, M. Sonnenschein, P. Poschlod, J. M. Van Groenendael, L. Klimeš, J. Klimešová, S. Klotz, G. M. Rusch, M. Hermy, D. Adriaens, G. Boedeltje, B. Bossuyt, A. Dannemann, P. Endels, L. Götzenberger, J. G. Hodgson, A.-K. Jackel, I. Kühn, D. Kunzmann, W. A. Ozinga, C. Römermann, M. Stadler, J. Schlegelmilch, H. J. Steendam, O. Tackenberg, B. Wilmann, J. H. C. Cornelissen, O. Eriksson, E. Garnier, B. Peco, The LEDA Traitbase: a database of life-history traits of the Northwest European flora. *J. Ecol.* **96**, 1266–1274 (2008). |
| Taseski *et al.*, 2019 | 01 June 2021 | G. M. Taseski, C. J. Beloe, R. V. Gallagher, J. Y. Chan, R. L. Dalrymple, W. K. Cornwell, A global growth-form database for 143,616 vascular plant species. *Ecology*. **53**, 2614 (2019). |

| | | |
|---|---|---|
| TRY | 09 May 2021 | J. Kattge, G. Bönisch, S. Díaz, S. Lavorel, I. C. Prentice, P. Leadley, S. Tautenhahn, G. D. A. Werner, T. Aakala, M. Abedi, *et al.*, TRY plant trait database – enhanced coverage and open access. *Glob. Chang. Biol.* **26**, 119–188 (2020). |
| RAINBIO | 06 June 2021 | G. Dauby, R. Zaiss, A. Blach-Overgaard, L. Catarino, T. Damen, V. Deblauwe, S. Dessein, J. Dransfield, V. Droissart, M. Duarte, H. Engledow, G. Fadeur, R. Figueira, R. Gereau, O. Hardy, D. Harris, J. Heij, S. Janssens, Y. Klomberg, T. Couvreur, RAINBIO: A mega-database of tropical African vascular plants distributions. *PhytoKeys*. **74**, 1–18 (2016). |
| Rice *et al.*, 2019 | 21 June 2021 | A. Rice, P. Šmarda, M. Novosolov, M. Drori, L. Glick, N. Sabath, S. Meiri, J. Belmaker, I. Mayrose, The global biogeography of polyploid plants. *Nat. Ecol. Evol.* **3**, 265–273 (2019). |
| USDA | 23 May 2021 | USDA, NRCS. 2022. The PLANTS Database (http://plants.usda.gov, 05/23/2021). National Plant Data Team, Greensboro, NC USA. |
